# Disinfecting eukaryotic reference genomes to improve taxonomic inference from ancient environmental metagenomic data

**DOI:** 10.1101/2025.03.19.644176

**Authors:** Nikolay Oskolkov, Chenyu Jin, Samantha López Clinton, Benjamin Guinet, Flore Wijnands, Ernst Johnson, Verena E. Kutschera, Cormac M. Kinsella, Peter D. Heintzman, Tom van der Valk

## Abstract

Ancient environmental DNA is increasingly essential for reconstructing past ecosystems, particularly when palaeontological and archaeological tissue remains are absent. Detecting ancient plant and animal DNA in environmental samples often relies on using extensive eukaryotic reference genome databases for profiling shotgun metagenomics data. However, microbial contamination in these references can introduce substantial biases in taxonomic assignments, especially given the typical low abundance of plant and animal DNA in such samples. In this study, we present a method for identifying bacterial and archaeal-like sequences in eukaryotic genomes and apply it to nearly 3,000 reference genomes from NCBI RefSeq and GenBank (vertebrates, invertebrates, plants) as well as the 1,323 PhyloNorway plant genome assemblies from herbarium material from northern high-latitude regions. Our analysis reveals microbial-like sequences in many eukaryotic reference genomes, which are most pronounced in the PhyloNorway dataset. We provide a detailed map of the microbial-like regions, including genomic coordinates and taxonomic annotations. This resource enables the masking of microbial-like regions during profiling analyses, thereby improving the reliability of ancient environmental metagenomic datasets for downstream analyses.

## Introduction

Ancient environmental DNA (aeDNA) is a tool for studying past ecosystems, especially in contexts where traditional archaeological and palaeontological tissue remains, such as bones and seeds, are absent [1–4]. It consists of genetic traces left by organisms in the environment, such as soils, sediments, ice, or other environmental samples, and allows for the reconstruction of past biodiversity and ecological communities to provide insight into species extinction, vegetation changes, and ecosystem responses to climatic shifts and anthropogenic impacts.

The often limited amount of DNA that can be isolated from ancient environmental samples imposes significant constraints on analytical methods. Coupled with the often low relative abundance of plant and animal DNA preserved in most environments, as compared to microbes, aeDNA analysis primarily relies on a reference-based approach for taxonomic profiling, which assumes similarity between the aeDNA query and the reference genome sequences. Therefore, robust aeDNA-derived community reconstructions are dependent on the accuracy of read identification by comparison to genomic reference databases. Consequently, both the quality of aeDNA data and the reference databases is crucial for reliable inferences. Microbial-like sequences within reference genomic databases, that are either derived from non-endogenous sources (contamination) or similar to highly-diverged taxa (evolutionarily conserved or convergent), can be a potential source of false-positive taxonomic identifications.

The existence of contaminant-like sequences is a pervasive issue in reference genomes with multiple examples reported in the literature [5–7]. For instance, contaminated reference sequences, such as the presence of hippopotamus-like sequence in the alpaca mitochondrial reference genome [8] and human sequences in parasitic worm genomes [9], have led to inaccurate inferences of evolutionary relationships [10], divergence times [8], and horizontal gene transfer events [11]. The inclusion of such eukaryotic reference genome contamination, most commonly originating from microbial or human sources, can occur at any stage throughout the genome assembly process [12].

Several analytical approaches have been proposed to address the issue of microbial contamination in reference genomes. For instance, Lu and Salzberg [13] suggested a computational method for masking erroneous sequences from draft genomes of eukaryotic pathogens. This is implemented by splitting the draft pathogenic references into pseudo-reads and filtering them using *k*-mer based Kraken classification [14, 15] and Bowtie2 alignment [16] against the human genome and National Center for Biotechnological Information Reference Sequence Database (NCBI RefSeq) microbial references. Conterminator is another program for contamination detection in the NCBI GenBank, RefSeq, and non-redundant (NR) reference databases proposed by Steinegger and Salzberg [17]. The program operates by an exhaustive all-against-all sequence comparison across kingdoms by splitting reference sequences into short segments, extracting their *k*- mers, grouping the *k*-mers, and then performing cross-kingdom alignments of the representative sequences in order to predict the contaminating sequences. This approach identified over 2,000,000 contaminated entries in the GenBank database [18]. Furthermore, the Physeter [19] and CheckM [20] tools have also been used to estimate contamination levels in NCBI RefSeq bacterial genomes. Lastly, ongoing efforts by NCBI, such as introducing the FCS-GX tool [21], which uses hashed *k*-mer matches and a curated reference database, in addition to more traditional VecScreen [22] and BLAST [23], are retroactively reducing the prevalence of contaminant-like sequences within the NCBI RefSeq and GenBank databases.

However, these efforts do not address the same problems in alternative databases, such as those comprising genome-wide data, e.g. PhyloNorway, PhyloAlps [24], or in legacy versions of the NCBI RefSeq database [25, 26], that are commonly used in workflows for large-scale metagenomics analysis (e.g. Kraken [14, 15]). Therefore, there is a need for a generic tool that identifies and removes contaminant-like sequences, particularly those similar to bacteria and archaea, from any genomic datasets that will be used as reference sequences for ancient environmental metagenomics analysis. In addition, although the microbial NCBI RefSeq is one of the largest available reference databases that has previously been used for estimating the amount of contamination in eukaryotic reference genomes [13, 17], the advent of the more diverse and comprehensive microbial Genome Taxonomy DataBase (GTDB) [27] allows for greater sensitivity in identifying regions in genome assemblies that are characterised by containing microbial-like sequences.

The aim of this study was therefore threefold. First, we developed a generic algorithm applicable to any eukaryotic reference genome in FASTA format, which outputs exact genomic coordinates of microbial-like sequences in BED-format. The coordinate file can then be used to mask eukaryotic reference genomes for various applications, including taxonomic profiling from ancient metagenomics data. Second, we sought higher identification accuracy for regions of microbial-like sequences by using the curated and non-redundant microbial GTDB database - the most comprehensive of its type at present - with the goal of minimising false-positive discoveries in ancient environmental metagenomics studies. Lastly, to allow for future investigation of the sources and mechanisms of contamination, we annotated and summarized each genomic region identified as microbial-like by the relative contribution of each microbial taxon.

To showcase our approach, we aligned microbial sequences from the GTDB database to six panels of eukaryotic reference databases and identified genomic regions that are similar to bacterial and archaeal sequences. We show that up to 70% of a taxon’s reference genome assembly can have shared similarity with bacteria and archaea (microbial-like). After masking microbial-like regions from the reference genomes, we re-analysed two empirical ancient metagenomic datasets and showed that some eukaryotic species detections can be entirely driven by alignments of reads to microbial-like sequences. We anticipate that masking reference genomes for microbial-like sequences will greatly reduce reference-genome-derived false-positive taxonomic assignments in ancient and modern environmental metagenomic studies.

## Methods

We selected 4,294 reference genomes of varying degrees of completeness and from a broad spectrum of taxonomic groups. This included (1) chromosome-level reference genome assemblies for 96 plants, (2) 114 invertebrates, and (3) 162 non-mammalian vertebrate species available from NCBI RefSeq, release 213; (4) 566 chromosome- and scaffold-level mammalian genome assemblies from NCBI GenBank, release 254 (if a species had multiple assemblies, we selected the one with highest N50 value); (5) all 2,033 chromosome- and scaffold-level arthropod reference genomes available in NCBI GenBank, release 256; and (6) 1,323 genome-skimmed contig-level plant assemblies from the PhyloNorway project (DataverseNO, V1) [28]. We individually constructed Bowtie2 [16] indices for all 4,294 reference genomes in the six genome groups.

Next, we fragmented all microbial (bacterial + archaea) reference genomes present in the GTDB dataset ([27]; release 214 from the 28th of April 2023) into 60 bp long segments using a sliding window with a 10 bp step. This resulted in a collection of 2.6*10^10^ nucleotide sequences representing microbial sequencing data (reads), which we refer to as “pseudo-reads” in this study. These reads were aligned to each indexed eukaryotic reference genome using Bowtie2, with up to 10 multi-mappers retained per read. The retention of multi-mappers ensured that multi-copy microbial-like regions from the same microbe were also detected. In our testing, we discovered that keeping multi-mappers greatly improved the detection sensitivity for microbial-like regions in the eukaryotic reference genomes, with this gain saturating after retaining approximately 10 multi-mapped positions (Supplementary Figure 1). We considered genomic regions covered by at least one microbial pseudo-read as microbial-like. We visually validated a set of the microbial-like regions using the Integrative Genomics Viewer (IGV) [29], and confirmed their coverage by microbial pseudo-reads (Supplementary Figure 2). For additional details about the alignment procedure, see Supplementary Material S1.

We then used *samtools depth* [30] and *bedtools merge* [31] to detect and extract the coordinates of regions in the eukaryotic reference genomes that were covered by microbial pseudo-reads in BED format (Table 1, which also includes data on the abundance of the most prevalent microbes in each identified genomic region). The breadth of coverage of microbial-like sequences was computed as the fraction of reference genome nucleotides covered at least once by microbial pseudo-reads. We used *samtools* [30] and custom bash and R scripts for annotating the reference genomes with the most abundant source microbial species. The annotation was done both genome-wide and for each individual microbial-like region in the BED file. In the latter case, only the top 5 most abundant microbial taxa were recorded. The entire workflow is schematically presented in Figure 1 (see also Data and Code Availability).

**Figure 1.**
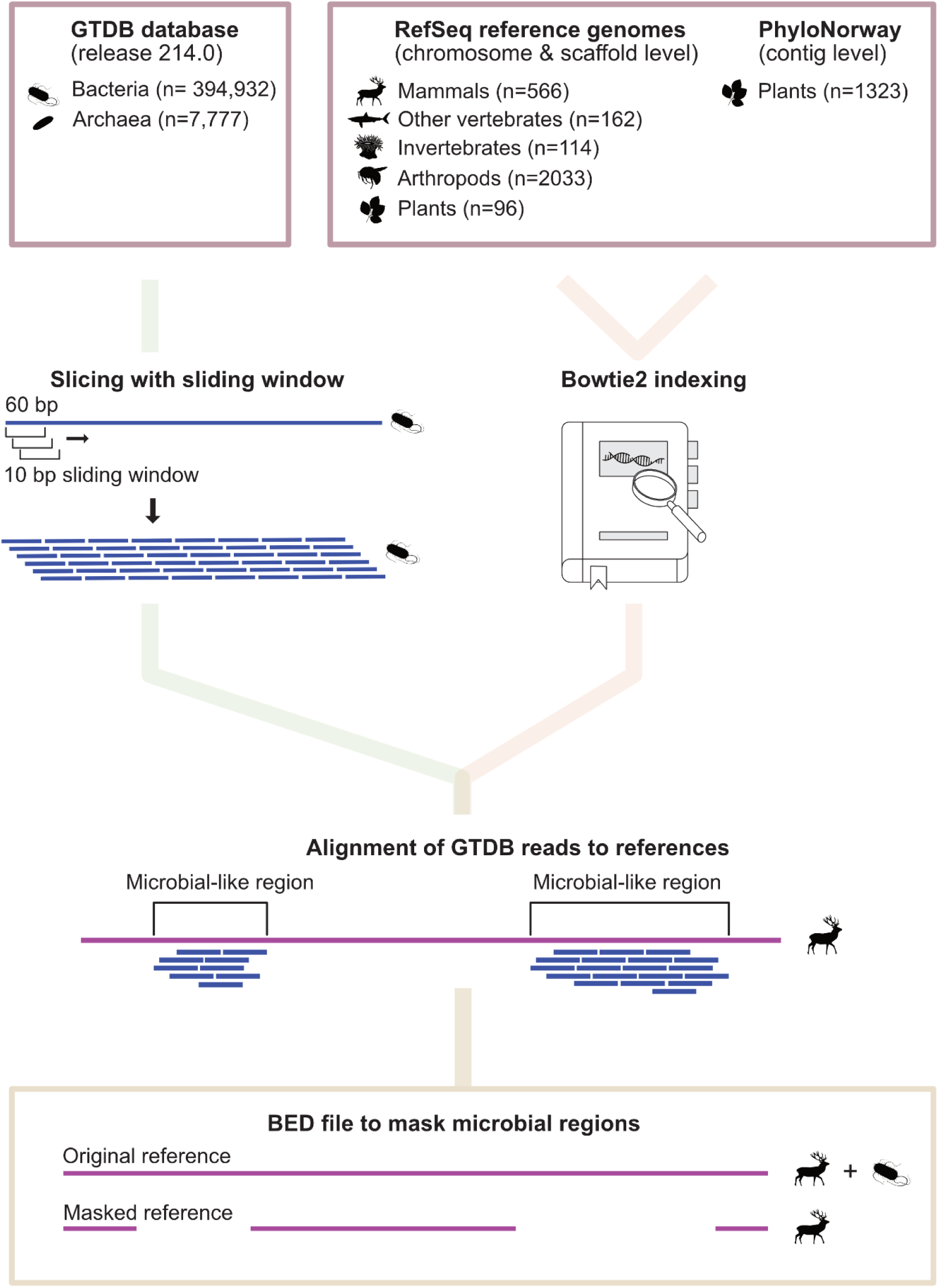
The workflow for detection of microbial-like sequences in eukaryotic reference genomes. Assemblies at different levels from different databases were subjected to the workflow. Microbial reference genomes from GTDB were fragmented into pseudo-reads (60 bp) and aligned to these genomes, retaining up to 10 multi-mappers to increase detection sensitivity. Contaminated regions were identified, annotated, and visualised, with microbial abundance summarised in BED files and validated using IGV.

**Table 1.**
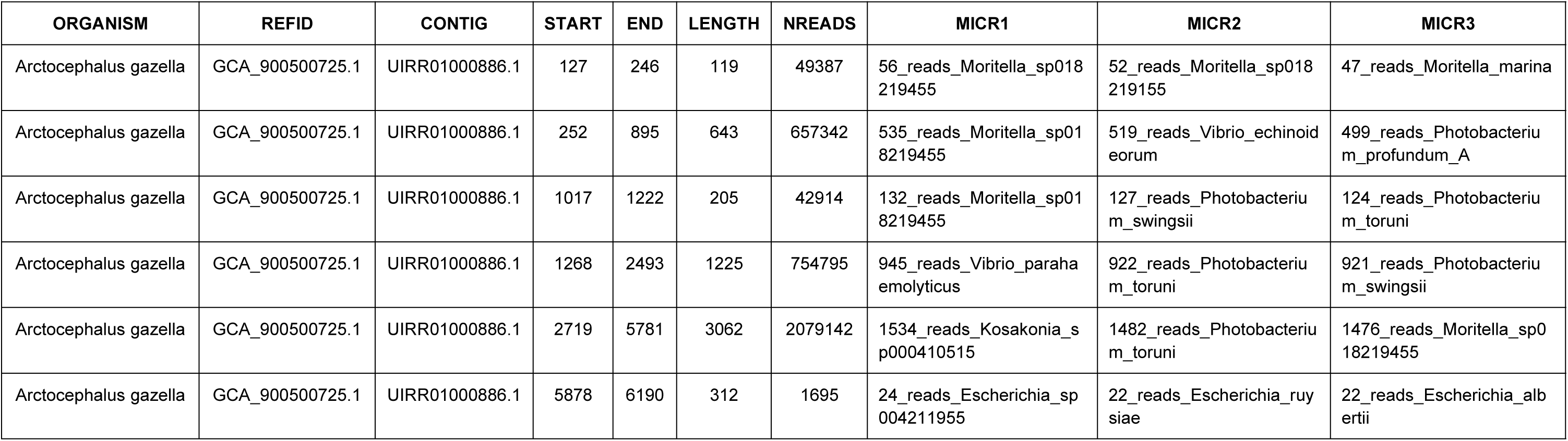
Example of BED-file with coordinates of microbial contamination of *Arctocephalus gazella*, reference genome GCA_900500725.1. The columns of the BED-file have the following notations: ORGANISM - scientific name of the organism, REFID - identification code of the reference genome, CONTIG - identification code of chromosome / scaffold / contig, START - start position of the segment of microbial contamination, END - end position of the segment of microbial contamination, LENGTH - length of the segment of microbial contamination, NREADS - number of pseudo-reads supporting the segment of microbial contamination, MICR1-3 - top 3 most abundant microbes contributing to the segment of microbial contamination; for example, the element “56_reads_Moritella_sp018219455” within MICR1 column denotes that *Moritella* sp018219455 was the most abundant microbe from that segment with 56 pseudo-reads.

The PhyloNorway dataset exhibited the highest proportions of microbial-like sequences of the genome groups. To validate our method, we therefore extracted the microbial-like (presumed exogenous) and remaining (presumed endogenous) segments from the PhyloNorway reference genomes with *bedtools getfasta* and *bedtools complement* [31] using their coordinates in the BED file. Next, we applied the Mash algorithm [32] (*mash dist* function was used) to construct a matrix of pairwise distances based on their *k*-mer composition among all species, separating the endogenous and exogenous segments. We then computed a Principal component Analysis (PCA) on the obtained matrix using the *scikit-learn* module in Python.

The workflow was verified against two empirical aeDNA datasets, which capture the flora and fauna from either across the Arctic or the Kap Kobenhavn Formation in Greenland [28, 33]. We used one sample from each study, i.e. cr9_67 from [28] (further referred to as the “Arctic sample”) and 69_B2_100_L0_KapK-12-1-35 [33] (further referred to as the “Greenland sample”). Adapter-removed reads from these samples were aligned with Bowtie2 [16] to the PhyloNorway reference genome assemblies, together with the Asian Elephant (EleMax1, GCF_024166365.1) and Human (GRCH38, GCF_000001405.40) reference genomes. These two latter mammalian references were added as decoys to attract mammalian reads via competitive mapping, since mammals were also reported in these samples in the original studies [28, 33]. We next applied *bedtools closest* [31] to compute the number of intersections of the aligned reads with the microbial-like sequences detected by our workflow in the PhyloNorway reference genomes. A custom R script was used to compute the null distribution of such intersections corresponding to random placement of the reads within the reference genomes.

## Results

After applying our workflow to a diverse set of eukaryotic reference genomes, we ranked the results by the percentage of the genome flagged as microbial-like sequence separately for each genome group (Figures 2 and 3, and Supplementary Tables 1-6).

**Figure 2.**
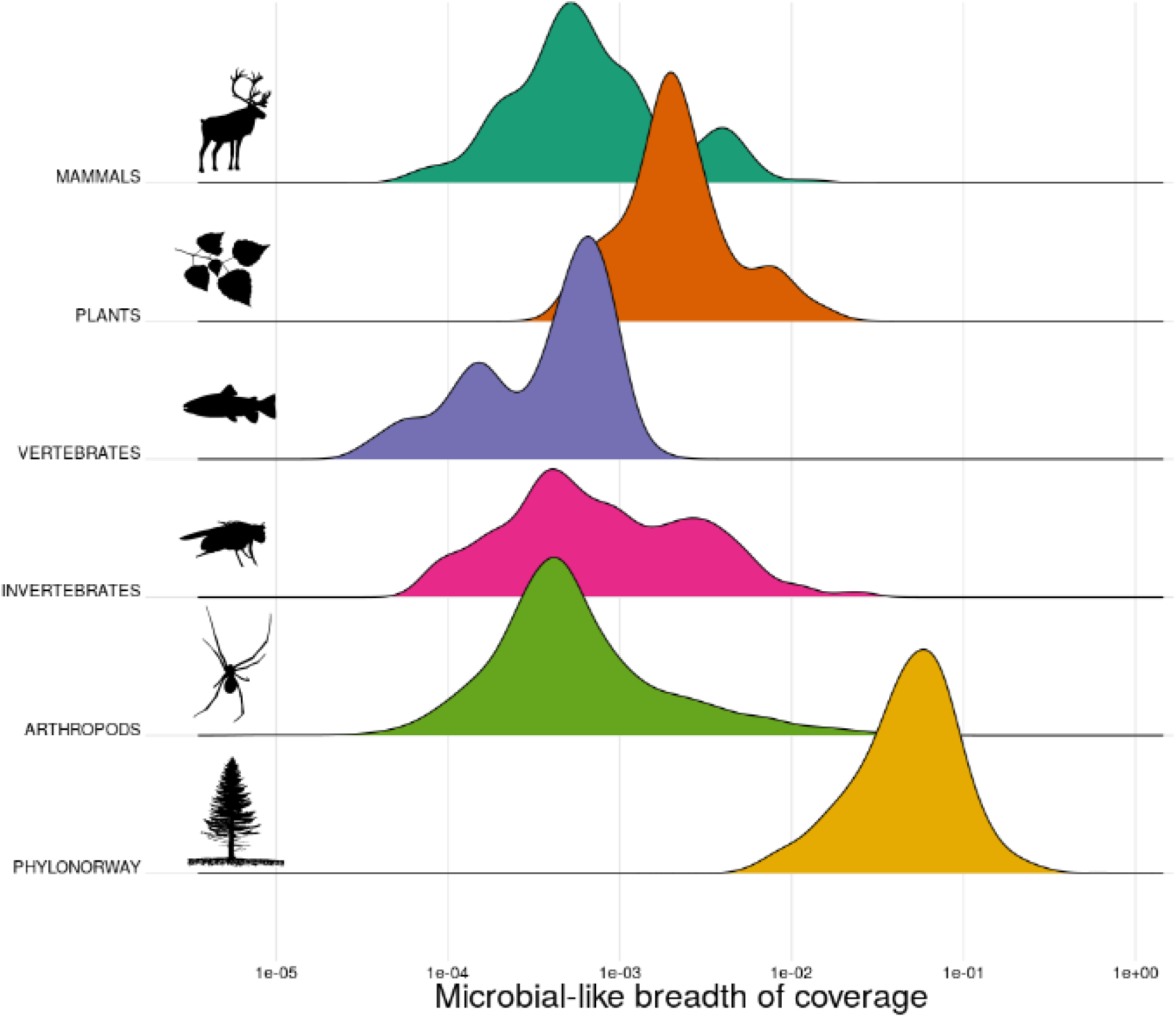
Distribution of microbial-like breadth of coverage (fraction of covered reference nucleotides) across the six reference genome groups from PhyloNorway or NCBI RefSeq. Mammalian genomes are represented by the genome with the highest contig N50 for each species sourced from the NCBI assembly database, which includes but is not limited to genomes from Refseq. The x-axis of the plot is on a log-scale. The y-axis represents the density estimates of the six datasets.

**Figure 3.**
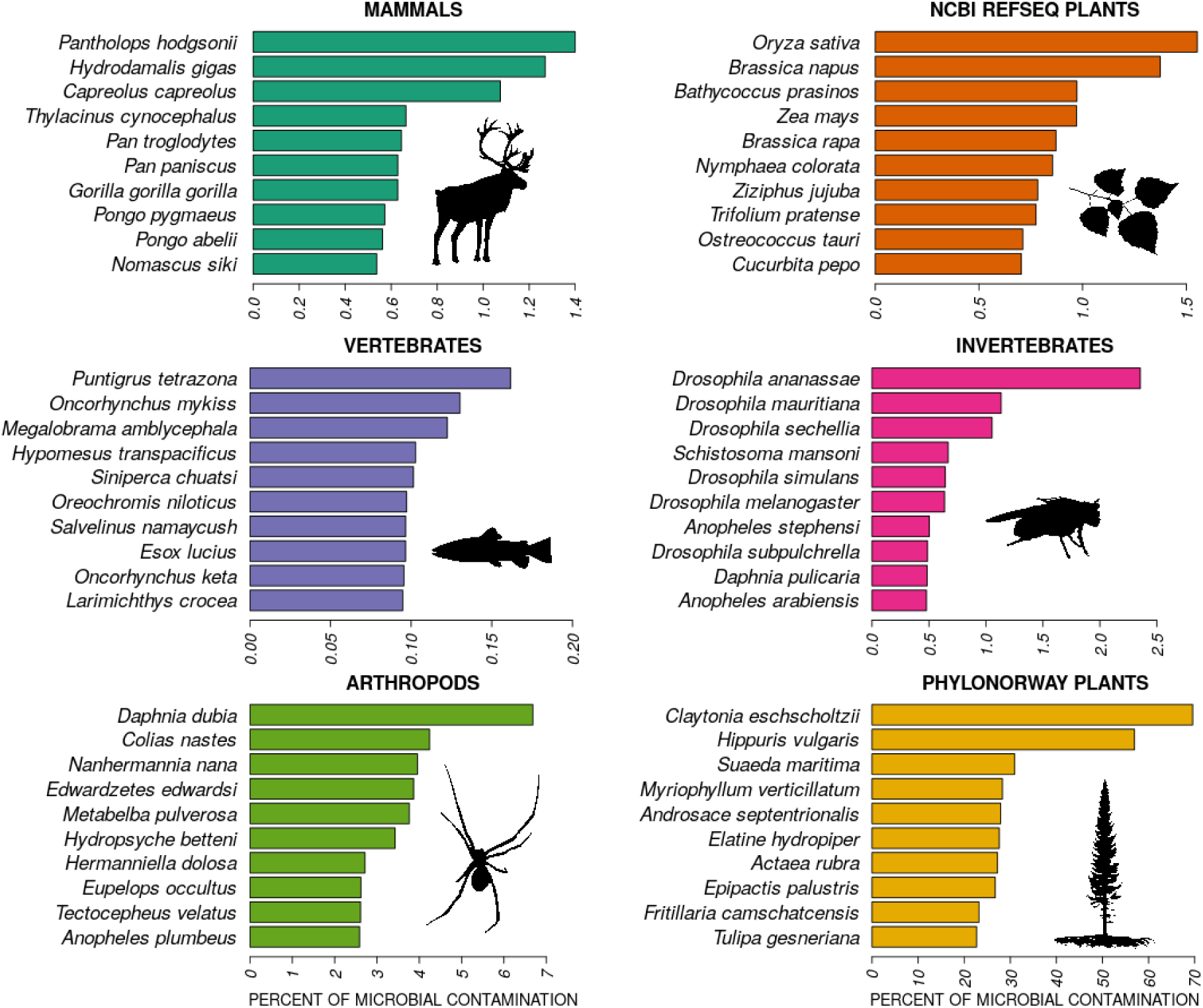
Reference genomes with the highest levels of microbial-like sequences for each genome group. Complete information is available in Supplementary Tables 1-6.

The non-mammalian vertebrate reference genomes exhibit the lowest overall levels of microbial-like sequence, i.e. <0.2% of the reference, compared to other genome groups, where the Tiger barb fish (*Puntigrus tetrazona*; NCBI id: GCF_018831695.1) has the greatest amount (0.16%). In contrast, mammals, plants, and invertebrate genomes often contained moderate degrees of microbial-like sequence, i.e. up to ∼1.5-2%, where Tibetan antelope (*Pantholops hodgsonii*; GCF_000400835.1; 1.4% microbial-like sequence), rice (*Oryza sativa*; GCF_001433935.1; 1.6%), and fruit-fly (*Drosophila ananassae*; GCA_017639315.2; 2.3%) contain the most microbial-like sequence in each respective group.

During the course of this study, NCBI RefSeq flagged the version of the Tibetan antelope reference genome used here (GCF_000400835.1) as containing a high magnitude of contamination and replaced it with an improved version (GCA_040182635.1). Using our workflow, we found that this reduced the percentage of microbial-like inserts in the Tibetan antelope genome from 1.4% to 0.12%, thereby indirectly validating the accuracy of our approach. Although the improved version of the Tibetan antelope reference genome contains an order of magnitude less microbial-like sequences, we suggest that the remaining microbial-like sequences detected here are likely due to the broader scope of the microbial dataset we used for the detections. The two reference genomes available for extinct organisms, Steller’s sea cow (*Hydrodamalis gigas*; GCA_013391785.1; 1.3%) and thylacine (*Thylacinus cynocephalus*; GCA_007646695.3; 0.7%), were found to be among the top five mammalian genomes with the most microbial-like sequence according to our method (Figure 3, Supplementary Table 1). We consider this plausible, as these genomes are derived from degraded samples with preservation conditions amenable to microbial contamination [34].

Among the mammalian genomes with the most microbial-like sequence, there is a significant over-representation of primates, consisting of 37 out of the top 45 mammalian genomes (i.e. 82%), while there are only 81 primate genomes out of the total 566 mammalian genomes assessed (i.e. 14%) (Fisher exact test, p=2.6*10^-11^). Second, bovids, including cattle (*Bos taurus*; GCA_947034695.1; 0.3%), wild yak (*Bos mutus*; GCA_027580195.1; 0.3%), and American bison (*Bison bison*; GCF_000754665.1; 0.2%), that are common organisms of interest in aeDNA studies, are placed among the top mammalian organisms with up to 9 Mb of their genomes consisting of microbial-like sequence.

Among plant reference genomes, rice (*Oryza sativa*; GCF_001433935.1; 1.6% microbial-like sequences), rapeseed (*Brassica napus*; GCF_020379485.1; 1.4%), corn (*Zea mays*; GCF_902167145.1; 1%) and pumpkin (*Cucurbita pepo*; GCF_002806865.1; 0.7%) have the highest fractions, i.e. 0.7-1.6%, of microbial-like inserts, corresponding to genomic lengths of 2-6 Mb (Figure 3, Supplementary Table 2). Interestingly, during the period of this study, the rice (*Oryza sativa*; GCF_001433935.1) reference genome, which we found to have the highest levels of microbial-like sequences, was suppressed by NCBI as a result of standard genome annotation processing, which can serve as an additional validation of our workflow. Invertebrates demonstrate similar levels, i.e. 0.5-2%, corresponding to genomic lengths of 1- 5 Mb, with the *Drosophila* genus among the invertebrates with most potentially contaminated reference genomes (Figure 3 and Supplementary Table 4).

GenBank arthropod reference genomes, which mostly comprise scaffold-level assemblies, on average demonstrate a comparable degree of microbial-like sequences as in NCBI RefSeq vertebrates and invertebrates (Figure 2). However, the most extreme examples show higher levels than those showcased from NCBI RefSeq vertebrates, invertebrates, and plants (Figure 3). For instance, the water flea (*Daphnia dubia*; GCA_013387435.1) has ∼7% of microbial-like sequences which corresponds to 7 Mb of genomic length), followed by the Labrador sulphur butterfly (*Colias nastes*; GCA_907164665.1; 4%; 20 Mb) (Figure 3 and Supplementary Table 5).

The PhyloNorway dataset, a collection of high-latitude skimmed plant genomes assembled from herbarium voucher specimens that is widely used in environmental ancient DNA studies [24, 28, 33], demonstrated particularly high levels of microbial-like sequences compared to all other datasets we analyzed in this work (Figure 2). For instance, the PhyloNorway genomes with the highest proportions of microbial-like sequences, such as grassleaf spring beauty flower (*Claytonia eschscholtzii*; 70%), common mare’s-tail plant (*Hippuris vulgaris*; 57%), and herbaceous seepweed (*Suaeda maritima*; 31%), were well above the levels observed in other datasets (Figure 3 and Supplementary Table 6).To further validate the difference in nucleotide composition between endogenous eukaryotic and microbial-like sequences in the PhyloNorway dataset, we visualized the two leading principal components of a PCA computed on their pairwise distances in Figure 4. We observed distinct clustering of microbial-like and endogenous regions, supporting the inference that the identified microbial-like sequences are not derived from plant genomes. In addition, reference sequences of *Hippuris vulgaris* projected on the hierarchical dendrogram built on pairwise *k*-mer distances between NCBI RefSeq plants and bacteria demonstrated that microbial-like sequences cluster together with bacterial genomes and endogenous sequences cluster with plant genomes (Supplementary Figure 3).

**Figure 4.**
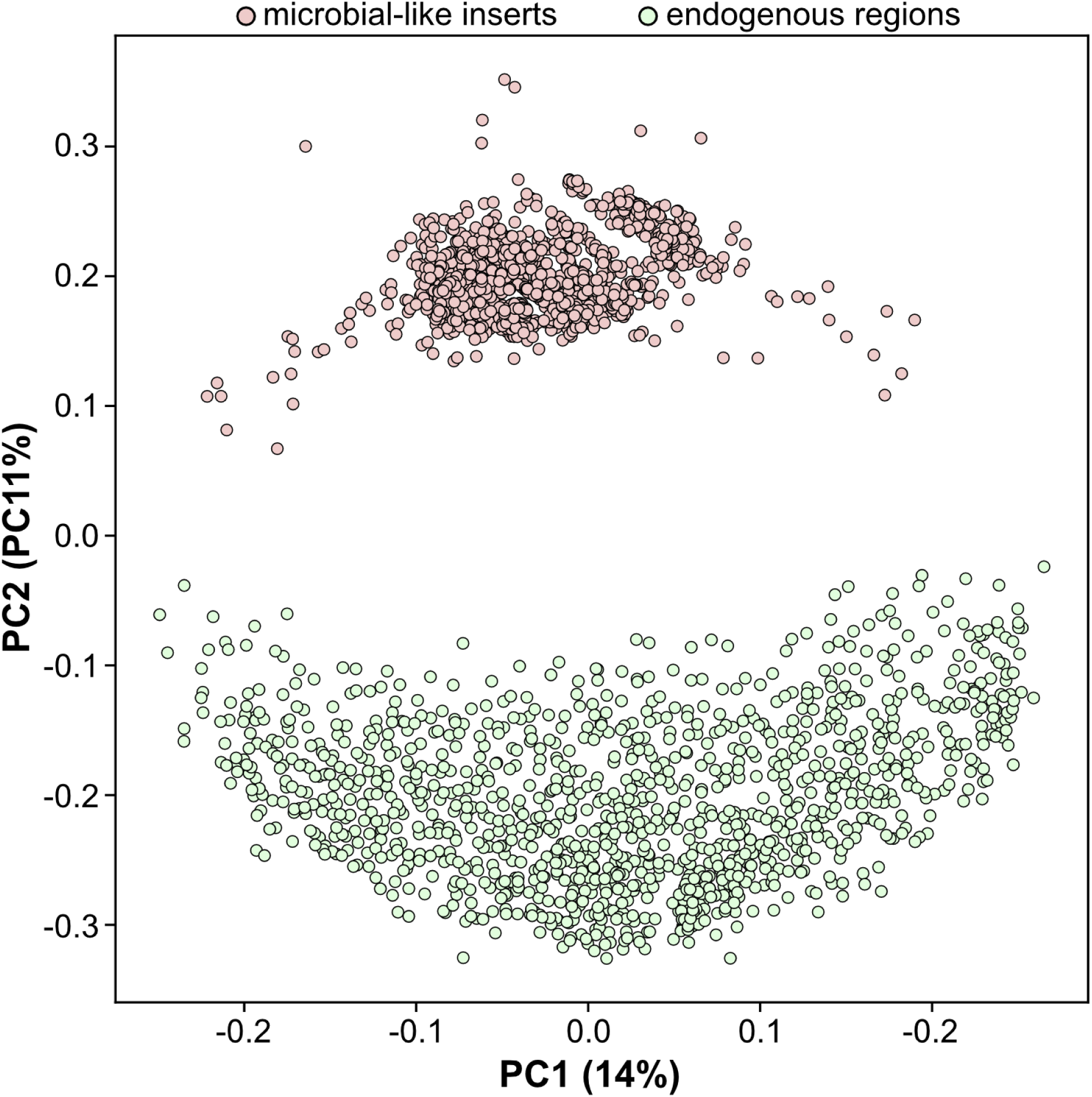
Principal Component Analysis (PCA) visualization of genomic pairwise distances between presumed endogenous (plant DNA) and exogenous (microbial-like) regions in the PhyloNorway dataset detected in this study. Each dot represents a single genome, with the light red dots representing regions identified as microbial-like and green dots as endogenous. The distinct clustering of endogenous and exogenous genomic segments suggests differentiation in their k-mer composition.

The aquatic plant genus *Hippuris* was found to be one of the most abundant in the two empirical studies examined and was reported from both northern Siberia (Arctic sample) [28] and Greenland (Greenland sample) [33]. Since this finding was based on alignments against the PhyloNorway reference genome assemblies, where *Hippuris vulgaris* was the only representative of *Hippuris* genus, and *Hippuris vulgaris* was shown by our analysis to be one of the species with the most extreme fractions of microbial-like sequences, we evaluated to what extent the conclusions of [28] and [33] could be affected by the presence of microbial-like sequences in the reference genome. The PhyloNorway reference genome assembly of *Hippuris vulgaris* consists of 433,631 contigs, which have a bimodal breath of coverage distribution for the microbial-like sequence fraction in our analysis, with modes at approximately 0 and 100% (Supplementary Figure 4). This indicates that a substantial proportion of *Hippuris vulgaris* contigs appear to be free from microbial-like sequences (the zero mode). In the Arctic sample however, a clear unimodal distribution of microbial-like fractions from the 20,213 *Hippuris vulgaris* contigs with at least one read mapped demonstrates that the vast majority of these contigs had close to 100% breath of coverage of microbial-like sequences (Supplementary Figure 5A). This implies that the *Hippuris*- identified reads from the Arctic sample have a much higher affinity to the microbial-like *Hippuris vulgaris* contigs, suggesting these reads originated from a microbial source. This indicates a potential mechanism for the discovery of *Hippuris* in [28]. In contrast, the reads attributed to *Hippuris vulgaris* in the Greenland sample from [33] mapped to 73,911 contigs that included both “endogenous” (to a larger extent) and “microbial-like” (to a lesser extent) contigs (Supplementary Figure 5B). Nevertheless, the peak at 100% of microbial-like fraction is not negligible, implying that the number of endogenous DNA sequences of *Hippuris vulgaris* in the Greenland sample was likely overestimated.

Of the 119,854 reads mapped in the Arctic sample, 116,483 (i.e. 97%) intersected with regions identified as microbial-like in the *Hippuris vulgaris* reference. To check whether this represents a statistically significant enrichment, we performed 300 random assignments of the 119,854 reads to the *Hippuris vulgaris* reference within the length limits of each contig, and demonstrated that approximately 58.8 ± 0.3 % would be a by-chance expectation if the intersection of mapped reads with the regions of microbial contamination was purely random. The observed 97% intersection is far beyond (p < 0.0033) the expected percentage (Supplementary Figure 6A). For the Greenland sample, where *Hippuris* was reported to be one of the most abundant genera in [33], 1,014,237 reads out of 1,367,627 reads, or 74%, mapped to microbial-like regions of the *Hippuris vulgaris* reference, which was again significantly higher (p < 0.0033) than the null expectation (Supplementary Figure 6B). For more details about *Hippuris vulgaris* follow up, see Supplementary Material S2. Therefore, for both the Arctic and Greenland samples, we conclude that the majority of their reads assigned to *Hippuris vulgaris* are of likely microbial origin.

We next sought to explore the potential mechanisms of the origins of microbial-like sequences in mammalian reference genomes. To achieve this, we quantified the abundance of the most common microbes in each eukaryotic reference genome and compared the reference genomes based on the patterns of microbial presence observed. The most common microbes across the mammalian reference genomes with the highest levels of microbial-like sequences form several clusters (Figure 5). First, the highly abundant *Streptococcus* sp000187445 bacterium is shared across six equid reference genomes (*Equus quagga burchellii*, GCA_026770645.1; *Equus przewalskii*, GCF_000696695.1; *Equus caballus*, GCF_002863925.1; *Equus asinus*, GCF_016077325.2; *Equus quagga*, GCF_021613505.1; *Equus asinus asinus*, GCA_003033725.1) and the white rhinoceros (*Ceratotherium simum simum*, GCA_023653735.1). Since these seven reference genomes were submitted to NCBI by different centers, lab contamination as a source for the microbial-like sequences is unlikely. The co-occurence of *Streptococcus* sp000187445 in equids and rhinos is intriguing, as these taxa comprise part of the odd-toed ungulates (order Perissodactyla). The remaining perissodactyl in the dataset, South American tapir (*Tapirus terrestris*), had the next highest abundance of *Streptococcus* sp000187445 but fell outside of the perissodactyl cluster. This suggests that *Streptococcus* sp000187445 could be a probiotic microbe endogenous to the perissodactyl microbiome or that part of the ancestral perissodactyl genome was evolutionarily convergent with *Streptococcus* sp000187445. Second, the D16-34 sp910588485 bacterium (belonging to genus *Adlercreutzia*) is highly abundant and shared by Snow sheep (*Ovis nivicola lydekkeri*, GCA_903231385.1) and Scimitar oryx (*Oryx dammah*, GCF_014754425.2) reference genomes produced by different centers. The two mammalian species belong to the Bovidae family which suggests some plausible similarity in their microbiomes or alternatively evolutionary convergence. Analogously, reference genomes, produced by different centres, of four mammalian species belonging to family Canidae, i.e. maned wolf (*Chrysocyon brachyurus*, GCA_028533335.1), arctic fox (*Vulpes lagopus*, GCF_018345385.1), dingo (*Canis lupus dingo*, GCF_003254725.2), and domestic dog (*Canis lupus familiaris*, GCF_013276365.1) share highly abundant microbial-like sequences from *Paracoccus denitrificans B*, which is a soil-associated bacterium not previously shown to be related to the canid microbiome. Therefore, evolutionary convergence can be a plausible explanation for co-occurence of *Paracoccus denitrificans B*-like sequences in the reference genomes of Canidae mammals. In addition, at least two more large clusters including broad groups of both mammalian and microbial organisms can be distinguished: 1) an ungulate cluster driven by intermediately abundant *Aureimonas A endophytica*, *Aliidongia dinghuensis*, *Mycobacterium malmesburyense*, *Anaerotardibacter muris*, *Muriiphilus lacisalsi*, and 2) a non-human primates cluster driven by moderately abundant *Streptomyces griseoincarnatus*, *Streptomyces kurssanovii*, *Chromatium weissei*, *Zobellia laminariae*, *Caproicibacter* sp900184925, *Streptomyces* sp020873915 and *Paeniglutamicibacter antarcticus*. The latter two clusters suggest that microbial-like sequences from multiple microbes contributed to reference genomes of evolutionarily related organisms possibly due to shared ecological environments and hence similarities of their microbiomes or evolutionary convergence. In contrast, there are a few clusters which likely point at some commonalities that are not strongly host-associated. For example, *Tumebacillus A avium* is shared at high abundance between Sunda flying lemur (*Galeopterus variegatus*, GCA_004027255.2) and Asian black bear (*Ursus thibetanus thibetanus*, GCA_009660055.1), which are not closely related species and the reference genomes were produced by different research institutes. Figure 5 also demonstrates that many microbial species, such as *Spirillospora cremea*, *Azonexus* sp016617495, D16-34 sp910588485, *Anaerotardibacter muris*, *Chromatium weissei* and *Aliidongia dinghuensis*, are moderately abundant across a wide range of distinct mammals. Since these microbes are also typical inhabitants of soil and aquatic environments, we hypothesize that they either represent environmental contamination which was incorporated during the sampling, sequencing and genome assembly process, or can also be due to evolutionary convergence. For discussion of microbial-like sequences composition within NCBI RefSeq / GenBank plants, invertebrates, non-mammalian vertebrates, arthropods, and PhyloNorway plants, please see Supplementary Material S3 and Supplementary Figures 7-11.

**Figure 5.**
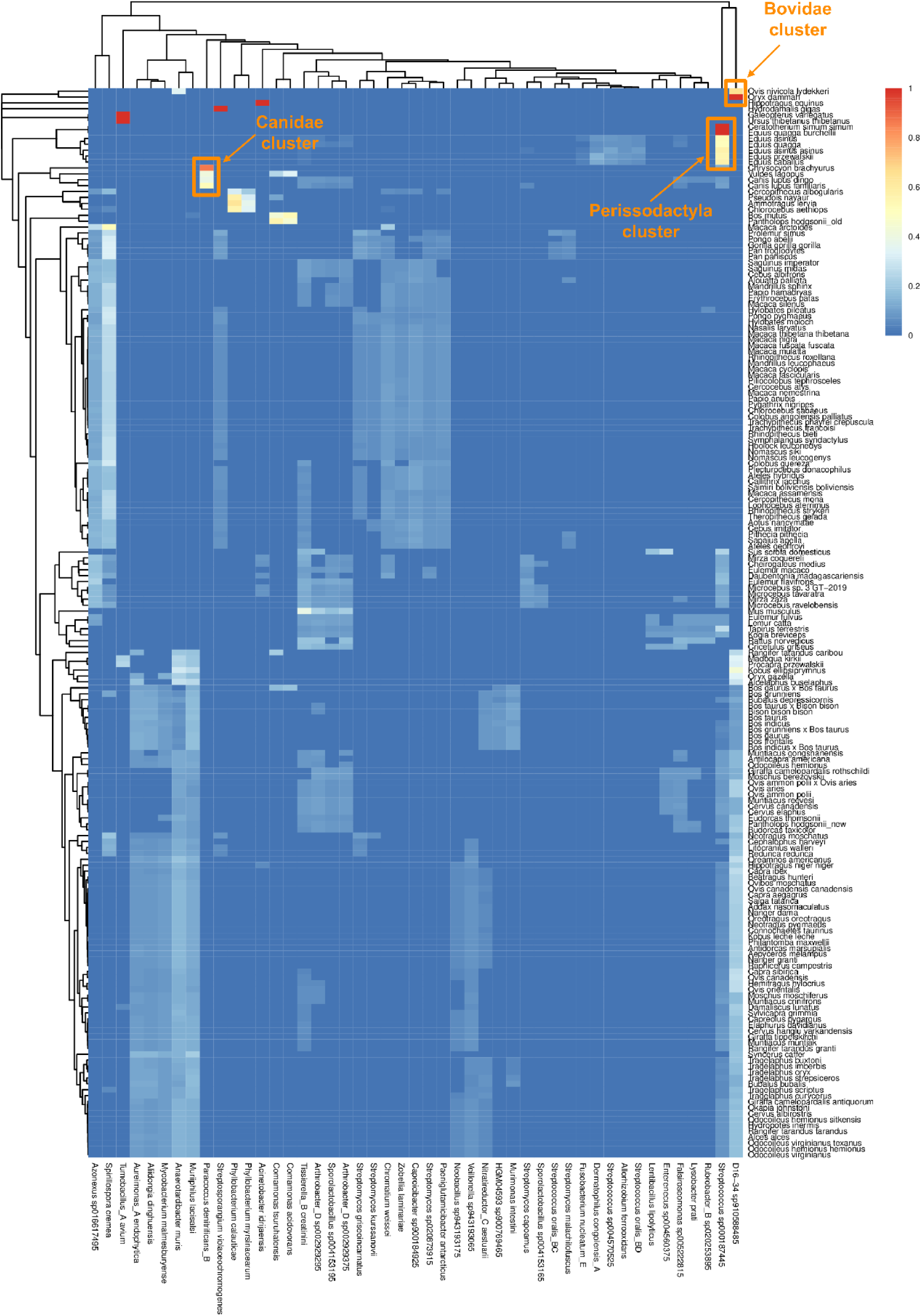
Abundance heatmap of microbial-like sequences across mammalian reference genomes. The columns represent microbial taxa contributing to the reference genomes of mammals displayed as rows. The color gradient indicates normalized abundance of microbial-like sequences (0-lowest,1- highest).

## Discussion

Microbial-like sequences present in reference genome databases represent a growing problem [35]. While human contamination was recognized some time ago to be one of the major challenges in ancient microbial genomics [9, 36], the opposite scenario of microbial contamination in animal and plant reference genomes became particularly evident in the rapidly developing ancient environmental DNA field [1–4], where reference-based organism discovery is widely used. Microbial contamination can occur at different steps of reference database generation [19, 20] and poses a serious risk of false-positive discovery, which, if neglected, can lead to erroneous results and interpretations in downstream analyses. Previous attempts to address this issue [12, 13, 17, 19, 20] have concentrated on flagging contaminated eukaryotic references without providing more comprehensive and quantitative information about specific locations and origins of microbial-like regions. Here we aim at mitigating microbial-like sequences with more precision and mechanistic understanding, while specifically concentrating on reference genomes that are particularly important for the area of ancient environmental DNA.

We present a novel method for detecting microbial presence within eukaryotic reference databases, and a collection of BED files (see example in Table 1) with coordinates of microbial-like sequences from a large custom dataset of ∼4,300 mammalian, non-mammalian vertebrate, invertebrate, arthropod, and plant reference genomes. The application of this method will allow researchers within the aeDNA field to mask portions of the genome with a potentially microbial origin. Therefore, rather than merely marking reference genomes as contaminated, our approach seeks to retrieve specific contigs and regions annotated with underlying microbial taxa. The method also enables more precise detections by utilizing the largest available microbial genome database (GTDB [27]), which includes both archaeal and bacterial genomes.

Although our approach follows a similar strategy suggested by Lu and Salzberg [13] and Steinegger and Salzberg [17], there are a few conceptual and technical differences. Lu and Salzberg [13] implemented splitting of eukaryotic reference genomes into pseudo-reads, screening them with Kraken [14, 15] and aligning them with Bowtie2 [16] against human and microbial references, while Steinegger and Salzberg [17] applied cross-kingdom *k*-mer matching across the NCBI RefSeq, GenBank, and NR databases. In contrast, we follow the opposite approach of splitting microbial (bacterial, archaeal) reference genomes into pseudo-reads and aligning them against eukaryotic references, which results in precise coordinates of microbial-like regions within eukaryotic reference genomes. The conceptual difference is that only eukaryotic pathogens were used in [13], while we utilise all NCBI RefSeq plant and animal references and the PhyloNorway dataset of plant genome assemblies. Therefore, our method is not specific to the NCBI databases but applicable to any custom nucleotide sequence in FASTA format. Another conceptual difference is that both [13] and [17] used the microbial NCBI RefSeq database, which has limited size and diversity compared to the non-redundant GTDB database [27], which we used in our testing (see Supplementary Material S4) and increases detection sensitivity to microbial-like sequences in eukaryotic reference genomes.

As microbial databases such as NCBI RefSeq and GTDB are continually updated with new assemblies, the masking of eukaryotic reference genomes with BED files from this study should not be considered an exhaustive solution. There will be a need to update the microbial-like regions presented here as microbial databases continue to grow. The importance of microbial database coverage can be seen from the study of Kjaer et al. [33], who used an older version of GTDB (release 95) as a decoy, in order to ensure that animal and plant hits were not originating from microbial reads. Nevertheless, we report in this study that a substantial amount of sequences attributed to the plant findings in the original work [33] are microbial-like. It is likely that a proportion of microbial-like reads were still remaining in [33] after filtering the data with the GTDB release 95, and further microbial-like sequence discovery became possible with the larger GTDB release 214 database used here.

There is currently no ultimate bioinformatic solution for distinguishing microbial contamination from true taxonomic hits in aeDNA studies. Here, we emphasize that we can only classify certain regions of eukaryotic reference genomes as “microbial-like”, which does not guarantee they are truly of microbial origin but could rather be due to sequence conservation or convergence, or from the potential insertion of microbial sequences into eukaryotic genomes.

Our study highlights the need to avoid using sequencing data mapped to publicly available genomes, without also accounting for microbial-like regions within the reference genome assemblies. When working with only a handful of reference genomes, it is possible to evaluate the contamination of each individually, either through bioinformatic methods or by consulting the methods used to construct each assembly. However, this quickly becomes infeasible in metagenomic studies, where data is often mapped against hundreds or thousands of different reference genomes. Mapping sequence reads against potentially contaminated reference genomes can lead to spurious detections of animal and plant organisms. We therefore suggest, as a preventive measure, to mask the microbial-like regions in the reference genomes before performing mapping, or adding a validation after mapping to confirm that the detection signal does not derive from the microbial-like regions.

## Acknowledgments

NO, CMK and VEK are financially supported by Knut and Alice Wallenberg Foundation as part of the National Bioinformatics Infrastructure Sweden at SciLifeLab. PDH and FW were supported by the Knut and Alice Wallenberg Foundation (KAW 2021.0048 [PDH, FW] and KAW 2022.0033 [PDH]). EJ is supported by the Swedish Research Council (VR 2020- 04808). TvdV, BG, CJ and SLC acknowledge support from the SciLifeLab and Wallenberg Data Driven Life Science Program [KAW 2020.0239].

## Data and Code Availability

NCBI RefSeq reference genomes, release 213 (from 23rd of July 2022), were obtained from https://ftp.ncbi.nlm.nih.gov/refseq/release/, and NCBI GenBank reference genomes were downloaded from https://ftp.ncbi.nih.gov/genomes/genbank/. GTDB dataset of microbial reference genomes (bacterial and archaeal), release 214, can be accessed at https://data.gtdb.ecogenomic.org/releases/release214/214.0/, and the PhyloNorway project DataverseNO V1 Nordic plant contig-level reference genomes are available at https://dataverse.no/dataset.xhtml;jsessionid=dfe334bbadc5fb9c5eab4332d568?persistentId=doi:10.18710/3CVQAG&version=DRAFT. The empirical datasets [28, 32] were obtained from EMBL-ENA under project accession PRJEB43822 and PRJEB55522, respectively. The adapter-removed reads for the Arctic sample were downloaded from the following address: ftp://ftp.sra.ebi.ac.uk/vol1/run/ERR645/ERR6458938/cr9_67.truncated.fastq.gz, and the adapter-removed reads for the Greenland sample were downloaded from the ftp-address: ftp://ftp.sra.ebi.ac.uk/vol1/run/ERR104/ERR10493316/69_B2_100_L0_KapK-12-1-35_Ext-12_Lib-12.pair1.truncated.gz. The BED-files with coordinates of microbial-like sequences for each group of eukaryotic organisms can be downloaded from the SciLifeLab Figshare repository https://doi.org/10.17044/scilifelab.28380476. The workflow together with the pre-built datasets of microbial pseudo-reads and other helping files is available at the SciLifeLab Figshare https://doi.org/10.17044/scilifelab.28491956. A comprehensive vignette covering workflow usage is available at https://github.com/NikolayOskolkov/MCWorkflow. Custom scripts used for performing the analysis and computing the figures for the manuscript are available at https://github.com/NikolayOskolkov/MCManuscript.

## Supplementary Material

### S1. Alignment of microbial pseudo-reads to eukaryotic reference genomes

To generate microbial pseudo-reads, we utilized 394,932 bacterial and 7,777 archaeal reference genome sequences from the Genome Taxonomy Database (GTDB) release 214 [27] downloaded from https://data.gtdb.ecogenomic.org/releases/release214/214.0/. Each of the microbial references was fragmented into 60 bp long segments using a sliding window with a 10 bp step. The 60 bp length of microbial pseudo-reads was chosen to provide sufficient specificity of matching to eukaryotic references as it is twice as long as the conventional 30 bp lower threshold of specificity across organisms in the tree of life [14, 15]. As a result, we generated a set of 2.6*10^10^ microbial pseudo-reads. The eukaryotic reference genomes were individually indexed with Bowtie2 [16] using the following command line:

*bowtie2-build --large-index reference_genome.fna.gz reference_genome.fna.gz --threads 20*

Afterwords, microbial pseudo-reads were aligned to each indexed eukaryotic reference genome, and the alignments were sorted and indexed using the following command lines:

*bowtie2 --large-index -f -k 10 -x reference_genome.fna.gz --end-to-end --threads 20 --very-sensitive -U microbial_reads.fna.gz | samtools view -bS -F 4 -h -@ 20 - | samtools sort -@ 20 - > MicrReads_aligned_to_reference_genome.bam*

*samtools index -c MicrReads_aligned_to_reference_genome.bam*

It is reasonable to assume that some microbial sequences can map to multiple loci in eukaryotic reference genomes. Therefore, in order to increase sensitivity of discovery of contaminated regions, we allowed up to 10 multi-mapping pseudo-reads to be kept in the alignments (the flag *-k 10* in the Bowtie2 command line above). For estimating the optimal number of multi-mappers to keep, we performed alignments of microbial pseudo-reads to Gray short-tailed opossum (*Monodelphis domestica*, GCA_027887165.1) and African elephant (*Loxodonta africana*, GCF_000001905.1) reference genomes while varying the maximum number of multi-mapped positions to retain for a read (0, 5, 10, 25, or 50 positions). We recorded the total number of both mapped reads and discovered regions of microbial-like sequences (Supplementary Figure 1). We observed that both sensitivity metrics for both organisms saturated at ∼5-10 multi-mappers. We therefore decided to allow up to 10 multi-mapping pseudo-reads to be kept when performing alignments.

The microbial-like regions were detected by computing the breadth of coverage (boc) from the alignments with *samtools depth* as

*samtools depth -g 0×100 -a MicrReads_aligned_to_reference_genome.bam > boc.txt*

Here, we used the *-g 0×100* flag to account for contributions from multi-mapping microbial pseudo-reads to the total coverage.

### S2. Following up the microbial-like and endogenous regions within *Hippuris vulgaris* reference genome assembly from the PhyloNorway dataset

We used the annotation file *PhyloNorwayContigs_acc2TaxaID.txt* provided together with the PhyloNorway dataset for retrieving 433,631 contig ids corresponding to the taxid = 39321 of *Hippuris vulgaris*. The corresponding reference sequences for each contig id were extracted with *seqtk subseq* function from the seqtk toolkit https://github.com/lh3/seqtk and saved as the *genome_39321.fna* FASTA-file using the following command lines:

*grep -w 39321 PhyloNorwayContigs_acc2TaxaID.txt | cut -f2 > accids_39321.txt seqtk subseq merged_PhyloNorway.fna accids_39321.txt > genome_39321.fna*

Further, after we have inferred the coordinates of microbial-like regions of *Hippuris vulgaris* with our method, and generated the *coords_micr_like_39321.bed* BED-file, which is available at SiLifeLab Figshare https://doi.org/10.17044/scilifelab.28380476, we proceeded with *bedtools getfasta* [31], and extracted the *Hippuris vulgaris* reference sequences corresponding to the microbial-like regions:

*bedtools getfasta -fi genome_39321.fna -bed coords_micr_like_39321.bed -fo microial_like_seqs_genome_39321.fna*

Next, we applied *bedtools complement* [31] and *bedtools getfasta* to group the remaining (presumed endogenous) reference sequences of *Hippuris vulgaris* in a separate FASTA-file:

*cut -f1,2 genome_39321.fna.fai > genome.fai*

*bedtools complement -i coords_micr_like_39321.bed -g genome.fai > endogenous_coords_genome_39321.bed*

*bedtools getfasta -fi genome_39321.fna -bed endogenous_coords_genome_39321.bed -fo endogenous_seqs_genome_39321.fna*

In order to explore whether the microbial-like reference sequences of *Hippuris vulgaris* cluster together with bacterial or plant reference genomes, we computed the *k*-mer pairwise distances with Mash [32] using 91 NCBI RefSeq plant and 100 random bacterial NCBI RefSeq reference genomes as well as the two additional *Hippuris vulgaris* FASTA-files corresponding to endogenous and microbial-like sequences. We performed hierarchical clustering with the *hclust* function in R using the Ward method (Supplementary Figure 3). We observed that the inferred microbial-like sequences of *Hippuris vulgaris* were clustering together with bacterial NCBI RefSeq reference genomes while endogenous sequences grouped with plant reference genomes.

Next, for each of 433,631 contigs of *Hippuris vulgaris* we computed the fraction of microbial-like sequences using the coordinates, *coords_micr_like_39321.bed*, of microbial-like regions. We plotted the histogram, Supplementary Figure 4, of microbial-like fraction with *plot_hist.R* available at https://github.com/NikolayOskolkov/MCManuscript.

After we have explored the microbial-like content of the *Hippuris vulgaris* reference genome assembly from the PhyloNorway dataset, we aimed at investigating how this could affect the read assignment in [28] and [33] studies reporting *Hippuris* prevalence at certain periods of history. We downloaded adapter-removed reads in the form of FASTQ-files corresponding to two samples from [28] (“Arctic sample”) and [33] (“Greenland sample”), where high *Hippuris* abundance was reported in the original studies:

*wget* ftp://ftp.sra.ebi.ac.uk/vol1/run/ERR645/ERR6458938/cr9_67.truncated.fastq.gz

*wget* ftp://ftp.sra.ebi.ac.uk/vol1/run/ERR104/ERR10493316/69_B2_100_L0_KapK-12-1-35_Ext-12_Lib-12.pair1.truncated.gz

Since both mammalian and plant organisms were reported for those two samples in the original studies [28] and [33], we implemented the competitive mapping approach to disentangle the mammalian and plant reads, and proceeded with the reads uniquely aligned to the *Hippuris vulgaris* reference. To perform the competitive mapping, we built Bowtie2 [16] index of the *Hippuris vulgaris* reference genome concatenated with Asian Elephant (EleMax1, GCF_024166365.1) and Human (GRCH38, GCF_000001405.40) reference genome. Next, we performed Bowtie2 alignment of the downloaded reads to the indexed composite reference, and extracted only the reads mapping uniquely to the *Hippuris vulgaris* reference genome:

*cat EleMax1.fna Human38.fna genome_39321.fna > EleMax_Human_Hippuris.fna bowtie2-build --large-index EleMax_Human_Hippuris.fna --threads 20 bowtie2 --large-index -x EleMax_Human_Hippuris.fna --end-to-end --very-sensitive --threads 20 -U cr9_67.truncated.fastq.gz | samtools view -bS -q 1 -h -@ 20 - | samtools sort -@ 20 - | samtools view -bS -q 1 -h -@ 20 $(cat accids_39321.txt | tr ‘\n’ ‘ ’) > cr9_67.truncated.fastq.gz.aligned_to_genome_39321.bam*

From the alignment BAM-file, we retrieved the ids of contigs with at least one read aligned, and using the BED-coordinates, *microbial_like_coords_genome_39321.bed*, of microbial-like regions for *Hippuris vulgaris*, we computed the fraction of microbial-like sequences corresponding to each contig with at least one aligned read (Supplementary Figure 5).

To understand how often the aligned reads overlap with the inferred microbial-like regions of *Hippuris vulgaris*, we extracted the coordinates of aligned reads with *bedtools bamtobed*

*bedtools bamtobed -i cr9_67.truncated.fastq.gz.aligned_to_genome_39321.bam > cr9_67.coords_aligned_reads.bed*

and calculated the number of intersections between the coordinates of the aligned reads and the coordinates of inferred microbial-like regions using *bedtools closest* and custom bash / awk command lines:

*bedtools closest -a cr9_67.coords_aligned_reads.bed -b coords_micr_like_39321.bed -d > cr9_67.coords_aligned_reads_annotated_with_closest_micr_like_region.bed*

*cut -f7 cr9_67.coords_aligned_reads_annotated_with_closest_micr_like_region.bed | awk ‘{if($1==0)print $0}’ | wc -l >> number_of_observed_intersects.txt*

We discovered that the vast majority of aligned reads, i.e. 116,483 out of 119,854 reads mapped in the Arctic sample (i.e. 97%) and 1,014,237 out of 1,367,627 reads (i.e. 74%) in the Greenland sample, intersected with the regions previously identified as microbial-like in the *Hippuris vulgaris* reference. To check whether this represents a significant enrichment compared to random read placement, we performed 300 random replacements of the aligned reads and every time counted the number of their intersects with the coordinates of microbial-like regions using a custom R script, please see the whole procedure in the R script *shuffle_reads.R* available at https://github.com/NikolayOskolkov/MCManuscript. We produced the Supplementary Figure 6 using the recorded numbers of intersects between the shuffled reads and microbial-like regions and plotted them with *plot_hist.R* script.

### S3. Microbial-like sequence composition of reference genomes from NCBI RefSeq plants, invertebrates, non-mammalian vertebrates, arthropods and PhyloNorway plants

We used samtools [30] and custom bash and R scripts for annotating the eukaryotic reference genomes with microbial taxonomic names corresponding to the most abundant microbial-like sequences. The most abundant (top 10 for each organism) microbes and eukaryotic references with the highest levels (top 200) of microbial-like regions were summarized via a heatmap computed by the *pheatmap* R package, demonstrating microbial co-occurrence in some groups of mammalian organisms (Figure 5). By analogy with the mammalian microbial-like sequences abundance heatmap, similar clustering patterns can be observed in microbial-like sequence composition of NCBI RefSeq plants, invertebrates, non-mammalian vertebrates, arthropods and PhyloNorway plants, shown respectively in Supplementary Figures 7-11.

For example, *Stenotrophomonas* sp003504055 is shared at high and moderately high abundance across two clusters comprising the fruit fly genus *Drosophila* (Supplementary Figure 8). Similarly, for non-mammalian vertebrate taxa, *Methylocystis* sp011058845 is highly abundant and shared across freshwater fishes such as northern pike (*Esox lucius*, GCF_011004845.1), lake whitefish (*Coregonus clupeaformis*, GCF_020615455.1), lake trout (*Salvelinus namaycush*, GCF_016432855.1), Atlantic salmon (*Salmo salar*, GCF_905237065.1), brown trout (*Salmo trutta*, GCF_901001165.1), chum salmon (*Oncorhynchus keta*, GCF_012931545.1), rainbow trout (*Oncorhynchus mykiss*, GCF_013265735.2), coho salmon (*Oncorhynchus kisutch*, GCF_002021735.2), sockeye salmon (*Oncorhynchus nerka*, GCF_006149115.2), pink salmon (*Oncorhynchus gorbuscha*, GCF_021184085.1) and chinook salmon (*Oncorhynchus tshawytscha*, GCF_018296145.1) (Supplementary Figure 9).

There are also a few clear clusters of arthropod reference genomes that share common microbial-like sequences. For instance, *Enterobacter* sp000493015 is commonly present among reference genomes of butterflies, moths and wasps such as Labrador sulphur (*Colias nastes*, GCA_907164665.1), Asiatic rice borer (*Chilo suppressalis*, GCA_902850365.2), parasitic wasp (*Cotesia vestalis*, GCA_000956155.1), and queen butterfly (*Danaus gilippus*, GCA_018231785.1), whereas *Sphingomonas* sp017418975 is prevalent and shared in reference genomes of soil and leaf associated arthropods such as beetle mite (*Nanhermannia comitalis*, GCA_034697665.1), oribatid mites (*Nothrus palustris*, GCA_034697745.1; *Malaconothrus monodactylus*, GCA_034697245.1), terrestrial cave isopod (*Haplophthalmus danicus*, GCA_034700045.1) and springtail (*Isotomurus plumosus*, GCA_034696705.1) (Supplementary Figure 10).

In contrast to the NCBI reference genomes, the PhyloNorway dataset does not demonstrate obvious commonalities in terms of co-occurrence of microbial-like sequences. Instead, there is at least one group of microbes including *JC017* sp004296775, *Solirubrobacter* sp003344625, *Frankia californiensis*, *Frankia* sp917627385, *Frankia meridionalis*, *Geodermatophilus endophyticus_A*, *Spirillospora cremea*, *Modestobacter lapidis*, *Geodermatophilus* sp019799925, *Streptomyces capoamus*, *SACZ01* sp023369685, *Ancylomarina* sp009669305, which is shared across nearly all plant genome assemblies in the PhyloNorway dataset (Supplementary Figure 11). This reflects, in our opinion, the common sample storage, processing, and sequencing routines used for generating these genome assemblies rather than shared ecological or evolutionary factors.

### S4. Discovering microbial-like regions with microbial RefSeq pseudo-reads

In addition to the microbial pseudo-reads produced from the GTDB database, which included only bacterial and archaeal reference genomes, we have also generated a set of 1.1*10^10^ nucleotide sequences using the NCBI RefSeq microbial database, release 213. The latter contained 39,760 microbial reference genomes including 28,044 bacteria, 11,220 viruses, 459 archaea, 33 fungi and 4 protozoa. The RefSeq microbial pseudo-reads were prepared in the same way as described in the Methods section. Despite the potential redundancy (e.g. some bacteria such as *Escherichia coli* may have multiple versions of a reference genome), the RefSeq microbial pseudo-reads may be useful for discovering viral-like sequences in eukaryotic reference genomes. This analysis can be used complementary to the detection of microbial-like sequences with the GTDB pseudo-reads within the main workflow. Both GTDB and RefSeq microbial pseudo-reads are publicly available together with the workflow files via the SciLifeLab Figshare https://doi.org/10.17044/scilifelab.28380476. We found that in most cases, either the coverage by GTDB and RefSeq pseudo-reads had good agreement (Supplementary Figure 12), or the GTDB pseudo-reads provided higher resolution of discovery of microbial-like sequences (Supplementary Figures 13 and 14). Nevertheless, viral-like regions within eukaryotic genomes can only be inferred using the RefSeq microbial pseudo-reads.

When using this workflow with RefSeq (viral) pseudo-reads, it is important to carefully assess genomic fragments classified as viral-like sequences, as they may not represent free-living viral contaminants, but rather endogenous viral elements (EVEs), which are “fossilised” viral sequences integrated into the host genome. Establishing EVEs is a challenging problem and requires careful analysis to confirm that these sequences are not of exogenous viral origin [37]. Our approach can be used for detecting only recent EVEs, as our workflow relies on a mapping tool that performs poorly with highly divergent DNA sequences [38], a common feature of EVEs. Homology-based methods therefore offer a more effective alternative for detecting distant viral relationships due to their greater flexibility and sensitivity [37, 39, 40].

## Supplementary Figures

**Supplementary Figure 1.**
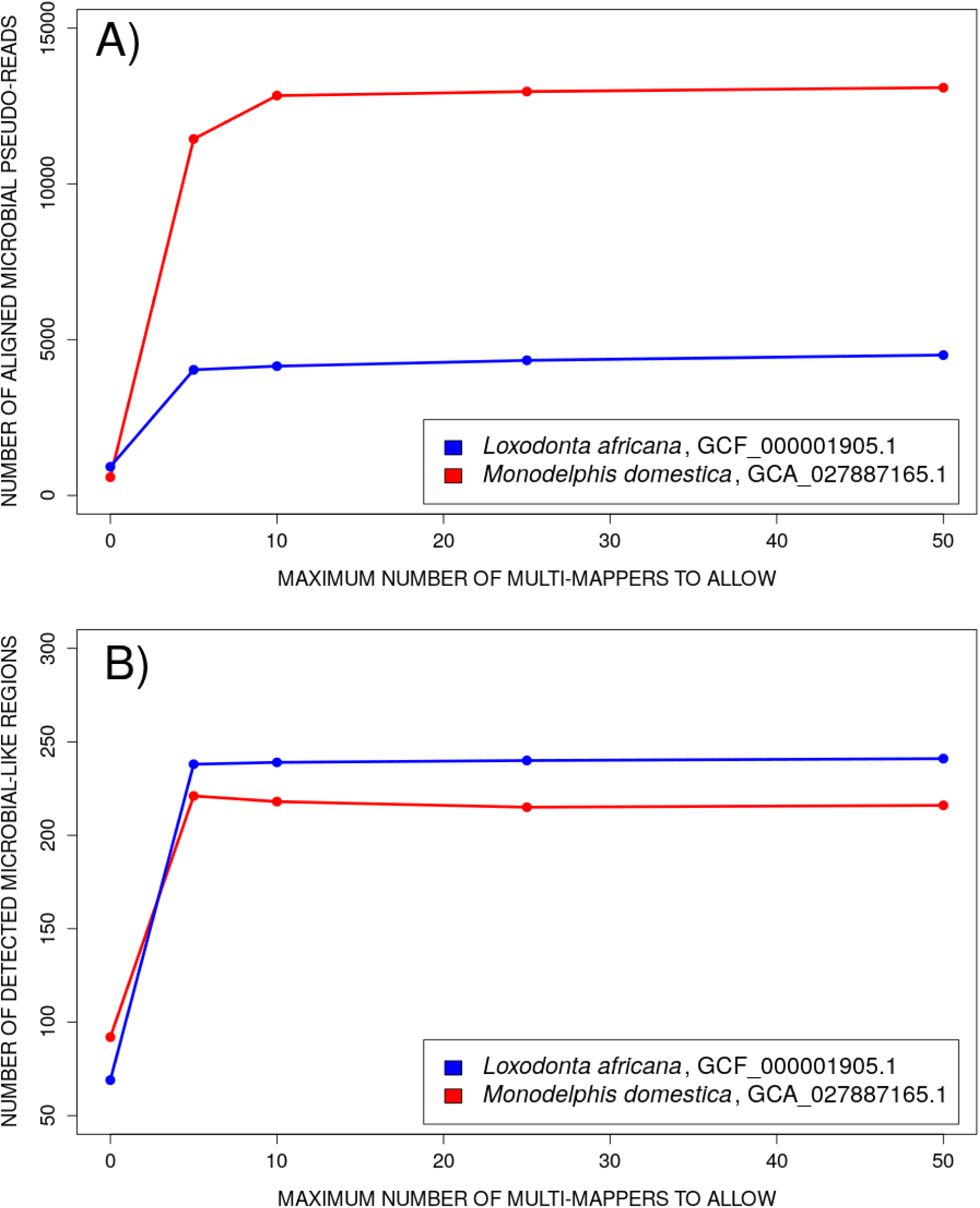
Sensitivity of discovery of microbial-like regions when aligning microbial pseudo-reads to Gray short-tailed opossum (*Monodelphis domestica*, GCA_027887165.1) and African elephant (*Loxodonta africana*, GCF_000001905.1) reference genomes with different numbers of multi-mapping pseudo-reads to retain.

**Supplementary Figure 2.**
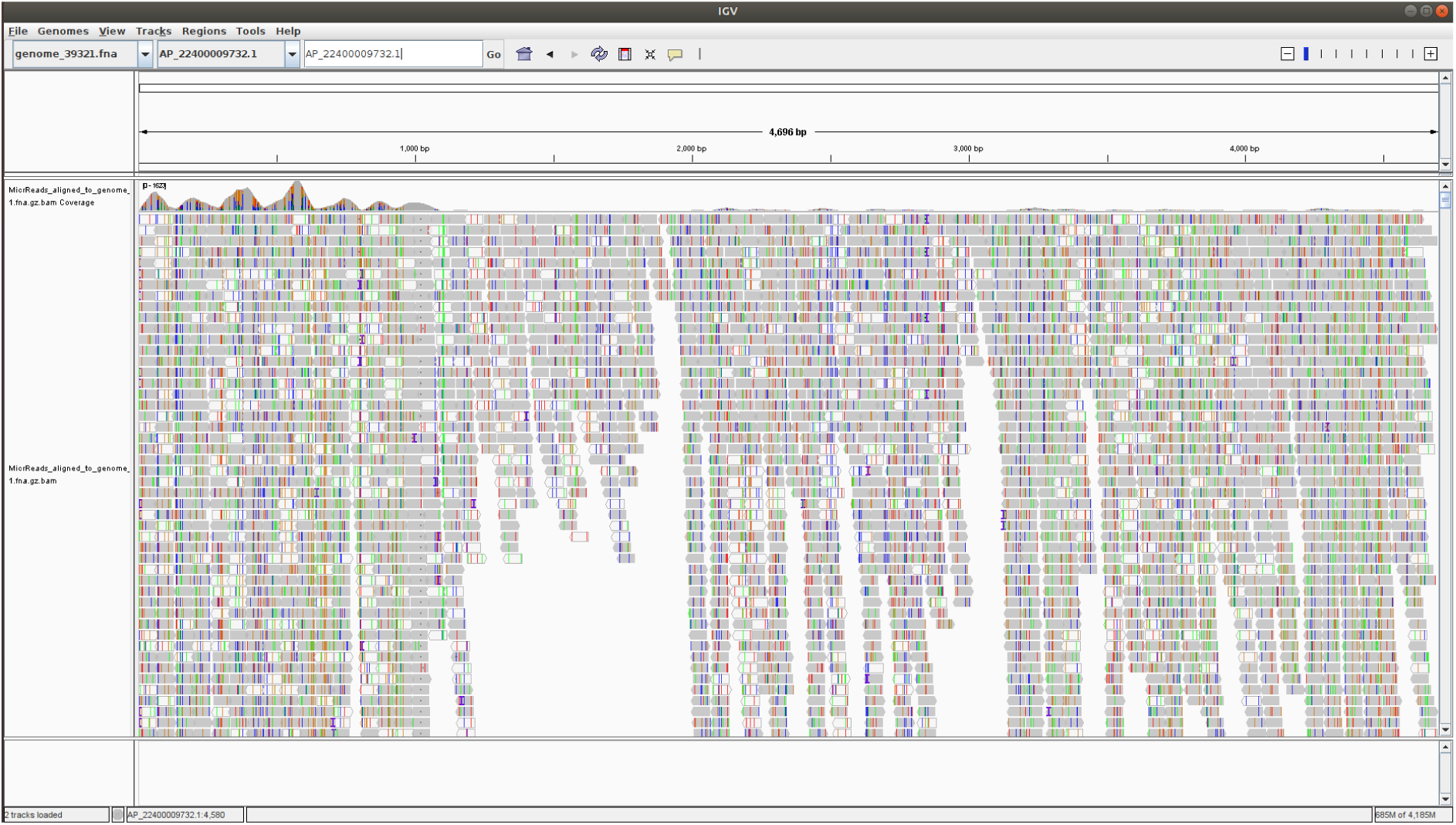
Example of coverage of detected exogenous regions by mapped bacterial pseudo-reads to the *Hippuris vulgaris* reference genome from the PhyloNorway dataset. The visualization is performed using the Integrative Genomics Viewer (IGV).

**Supplementary Figure 3.**
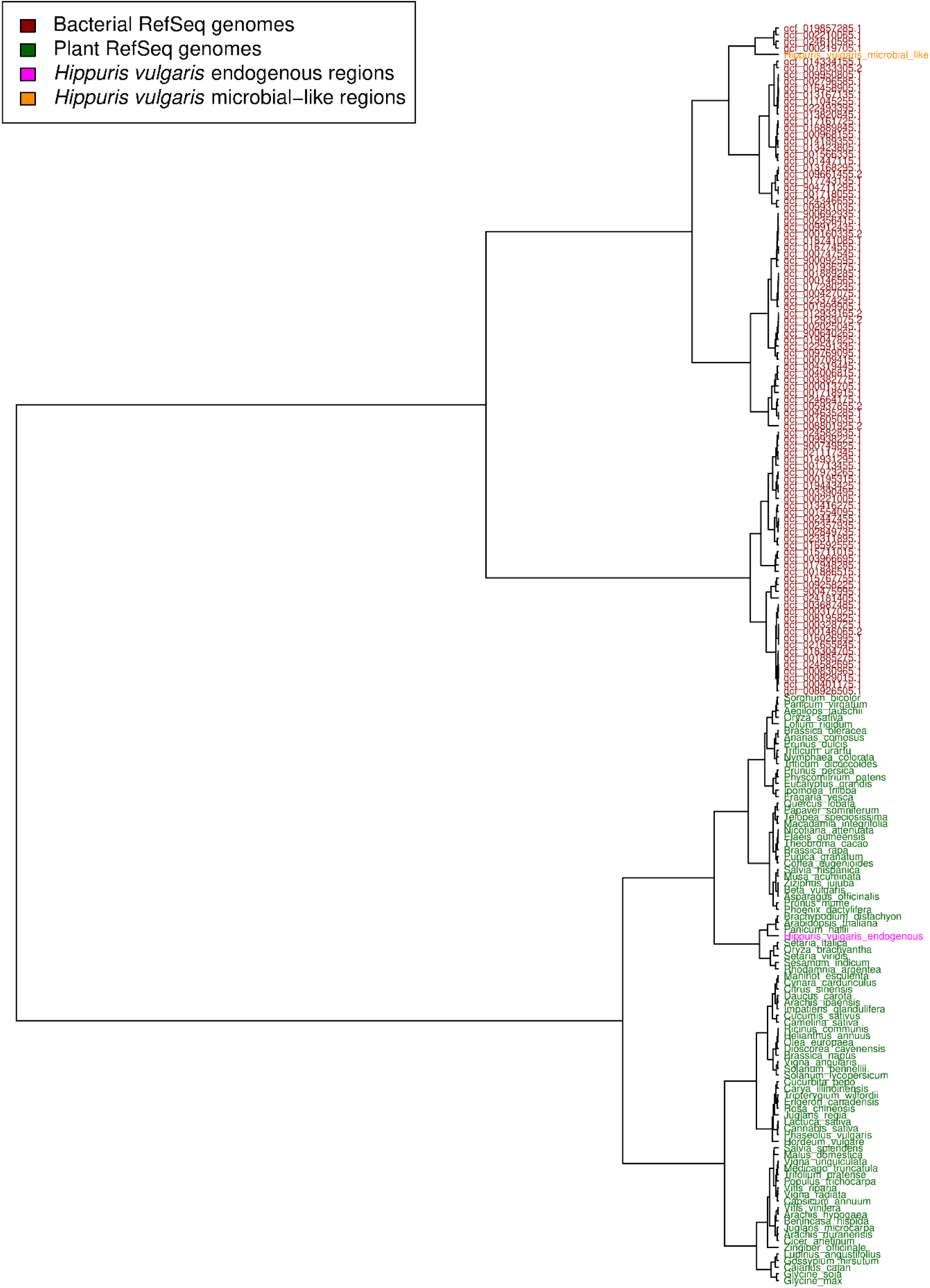
*Hippuris vulgaris* microbial-like (presumed exogenous) and remaining (presumed endogenous) segments from the PhyloNorway dataset projected on the hierarchical clustering dendrogram of NCBI RefSeq plants and bacteria computed using Mash [32] pairwise distances based on the *k*-mer composition of their reference genomes.

**Supplementary Figure 4.**
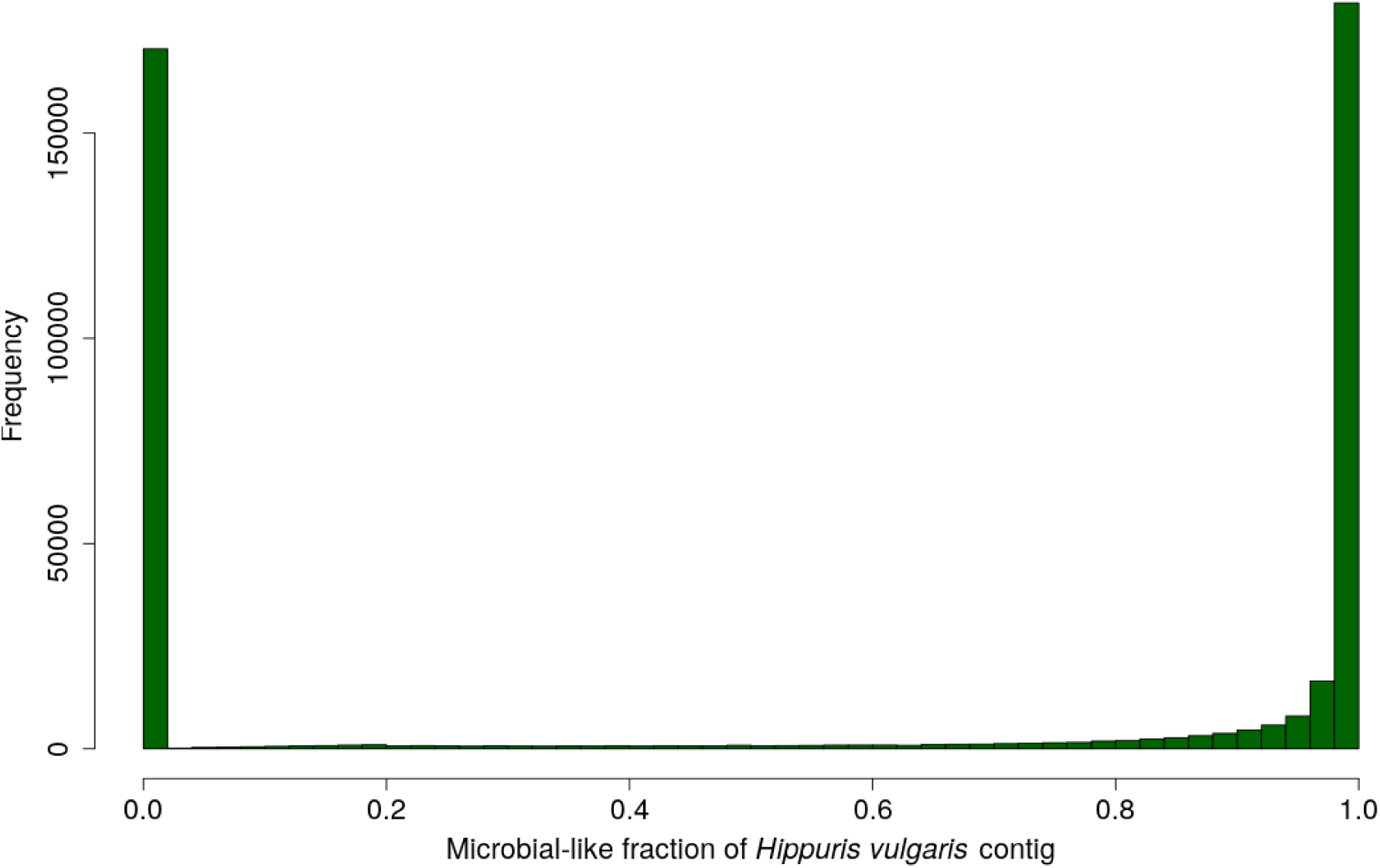
Distribution of microbial-like fractions of 433,631 contigs of *Hippuris vulgaris* from the PhyloNorway dataset profiled for microbial contamination in our analysis.

**Supplementary Figure 5.**
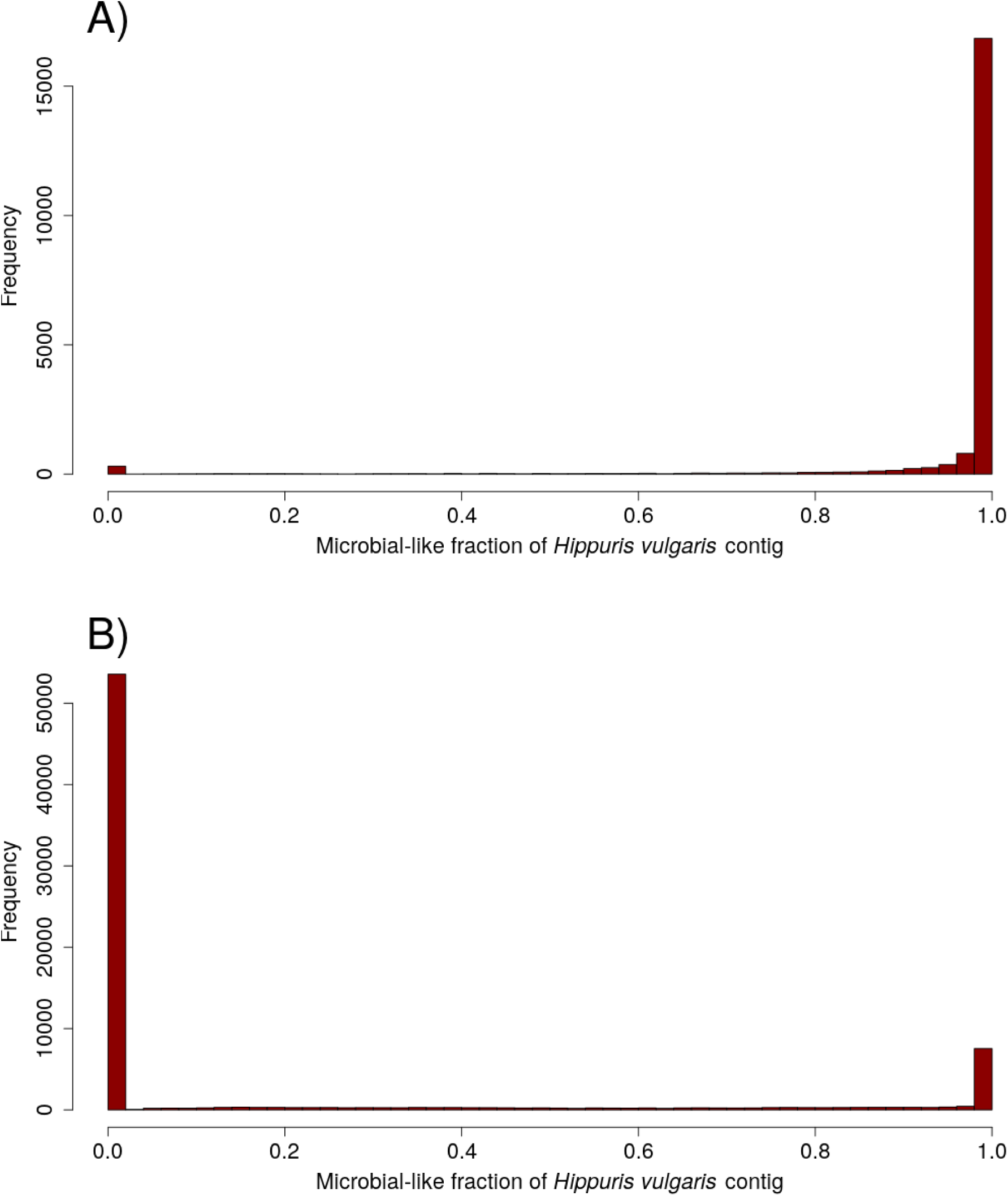
Distribution of microbial-like fractions of *Hippuris vulgaris* contigs with aligned reads for: A) Arctic sample cr9_67 [28] (20,213 contigs), and B) Greenland sample 69_B2_100_L0_KapK-12-1-35 [33] (73,911 contigs).

**Supplementary Figure 6.**
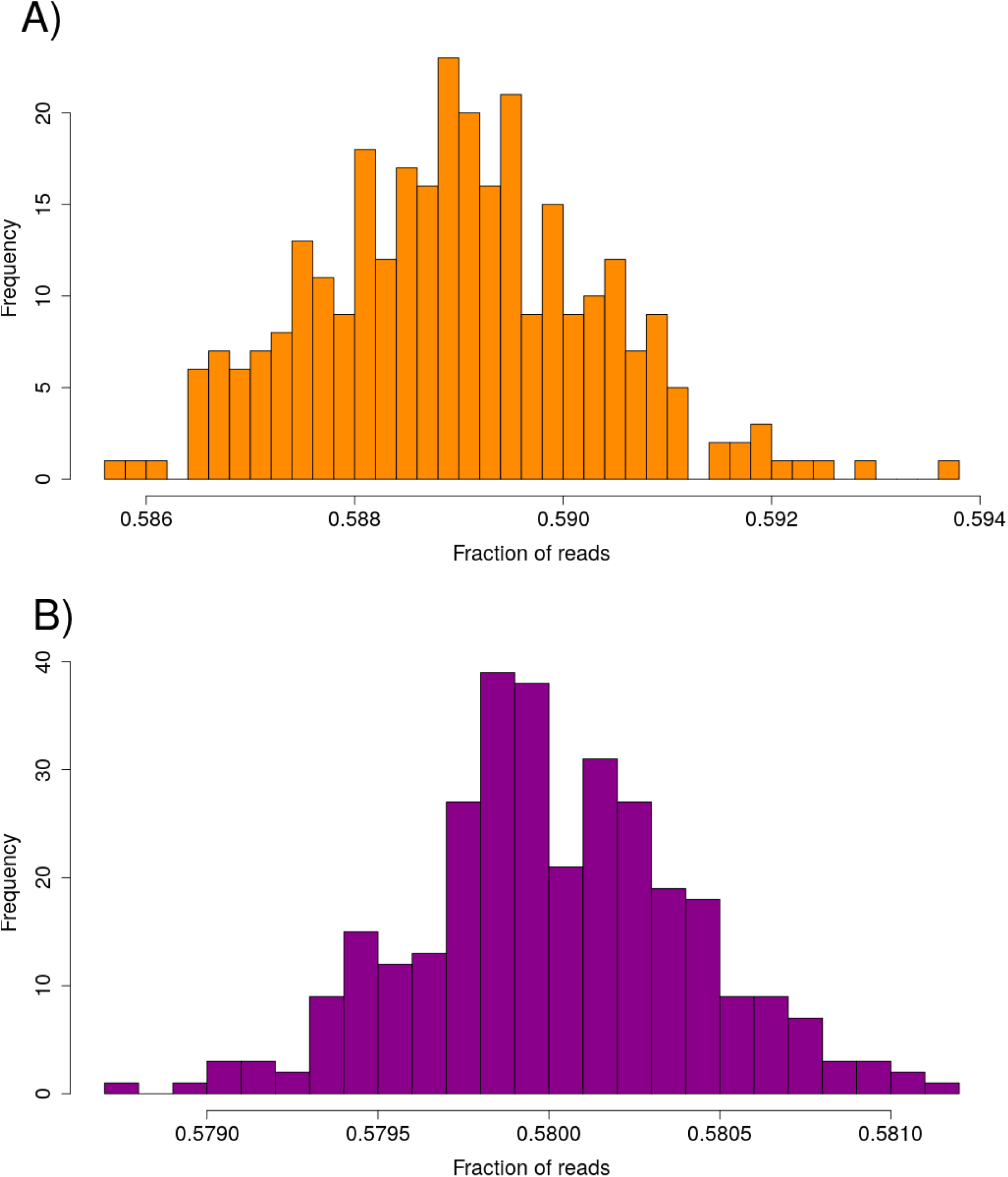
Verification of *Hippuris* hit from [28] and [33]. Intersection fraction of randomly assigned reads from: A) the Arctic sample cr9_67 [28], and B) the Greenland sample 69_B2_100_L0_KapK-12-1-35 [33], with microbial-like regions in the *Hippuris vulgaris* reference genome.

**Supplementary Figure 7.**
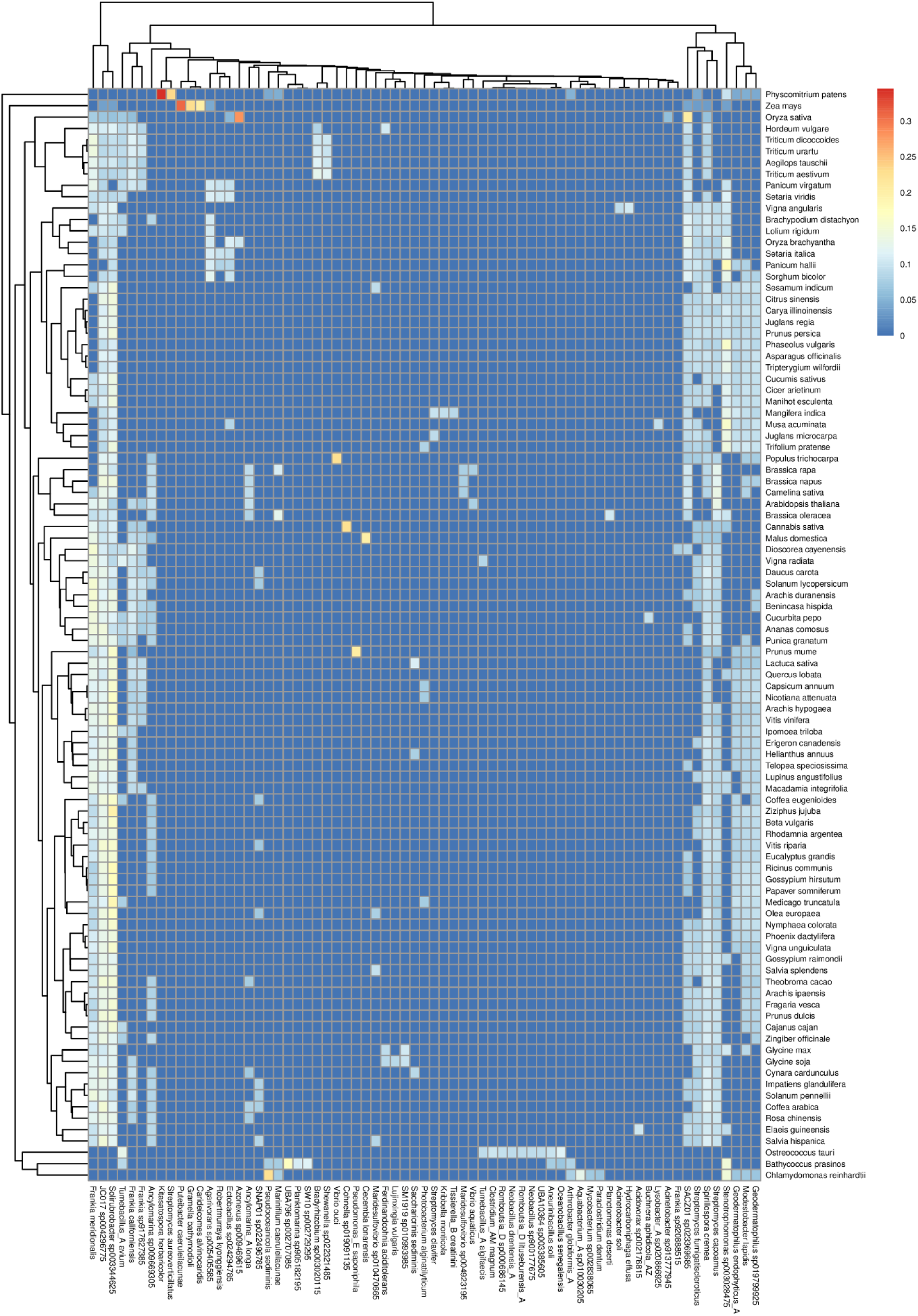
Abundance heatmap of microbial-like sequences across NCBI RefSeq plants. The columns represent microbial taxa contributing to the reference genomes of plants displayed as rows. The color gradient indicates normalized abundance of microbial-like sequences (0- lowest, 1-highest).

**Supplementary Figure 8.**
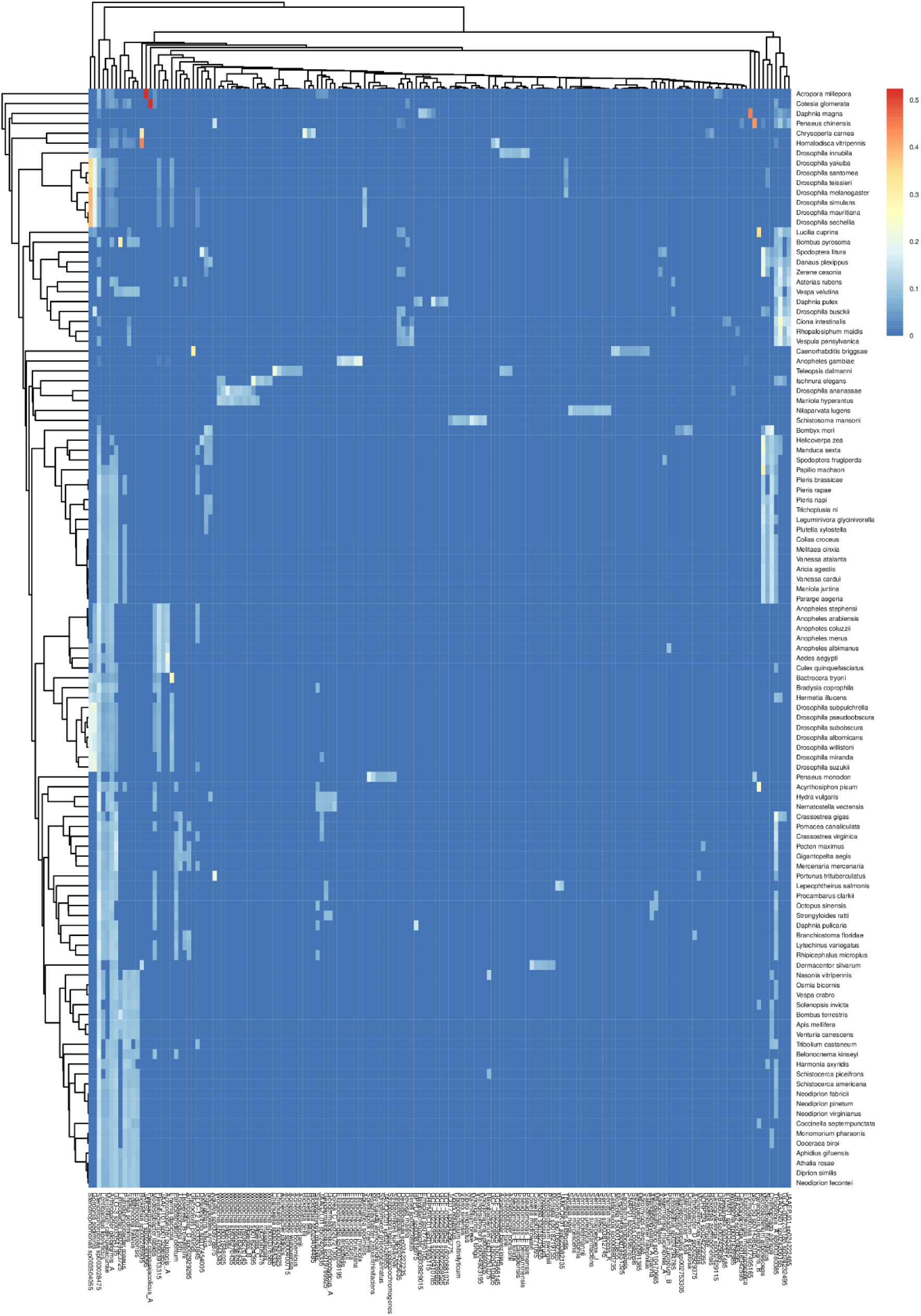
Abundance heatmap of microbial-like sequences across NCBI RefSeq invertebrates. The columns represent microbial taxa contributing to the reference genomes of invertebrates displayed as rows. The color gradient indicates normalized abundance of microbial-like sequences (0-lowest, 1-highest).

**Supplementary Figure 9.**
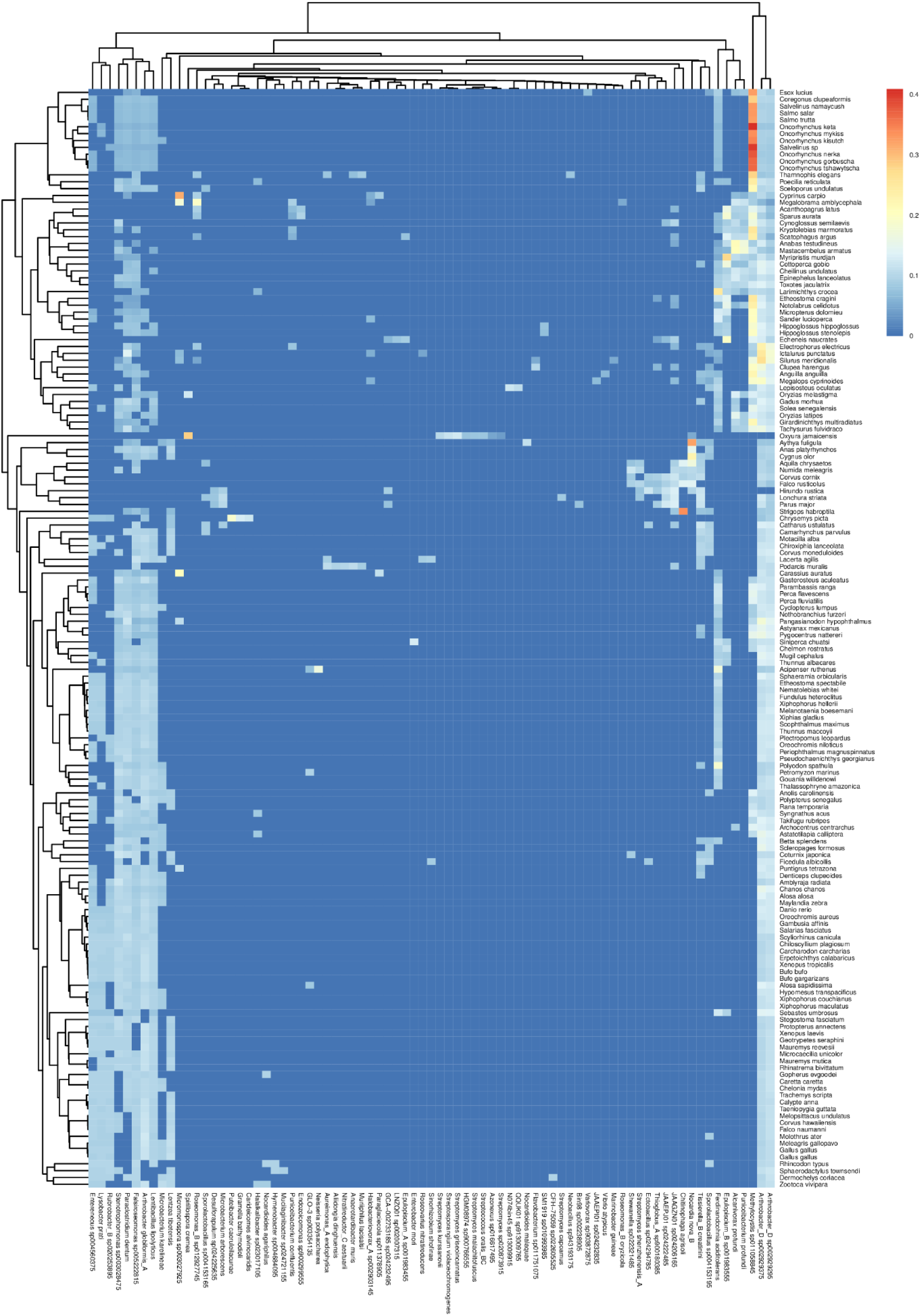
Abundance heatmap of microbial-like sequences across NCBI RefSeq non-mammalian vertebrate taxa. The columns represent microbial taxa contributing to the reference genomes of vertebrates displayed as rows. The color gradient indicates normalized abundance of microbial-like sequences (0-lowest, 1-highest).

**Supplementary Figure 10.**
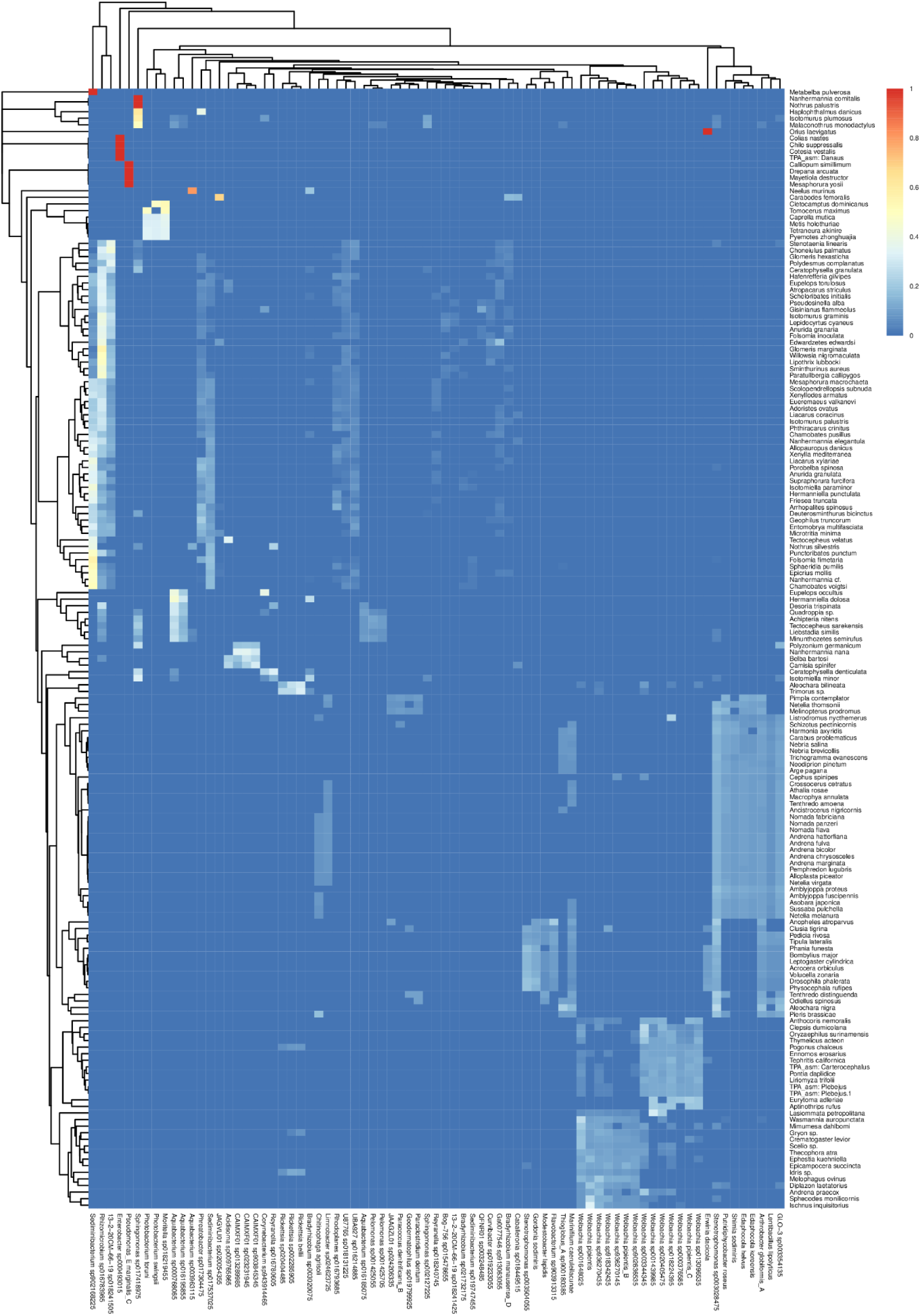
Abundance heatmap of microbial-like sequences across NCBI RefSeq arthropods. The columns represent microbial taxa contributing to the reference genomes of arthropods displayed as rows. The color gradient indicates normalized abundance of microbial-like sequences (0-lowest, 1-highest).

**Supplementary Figure 11.**
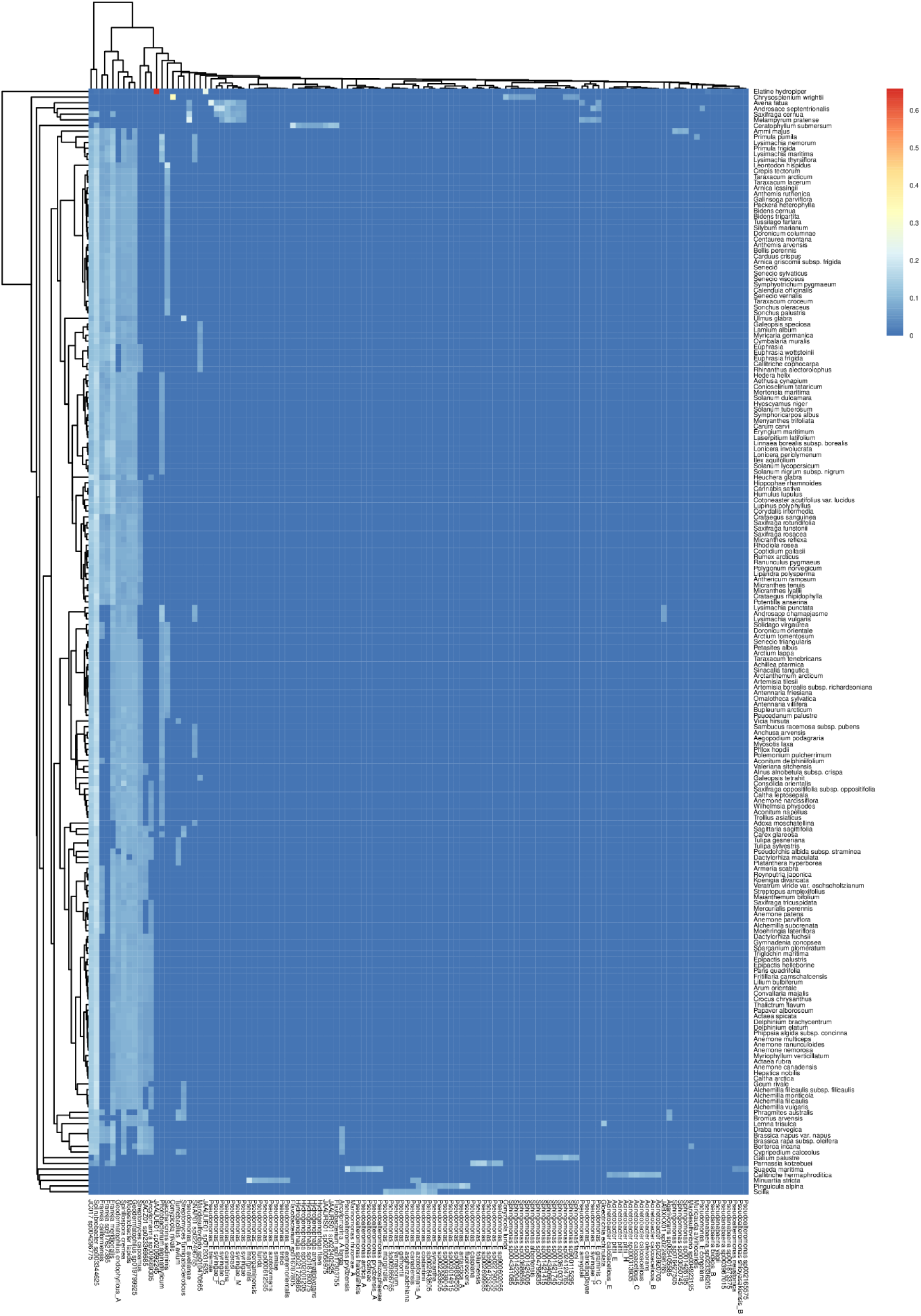
Abundance heatmap of microbial-like sequences across PhyloNorway plants. The columns represent microbial taxa contributing to the reference genomes of PhyloNorway plants displayed as rows. The color gradient indicates normalized abundance of microbial-like sequences (0-lowest, 1-highest).

**Supplementary Figure 12.**
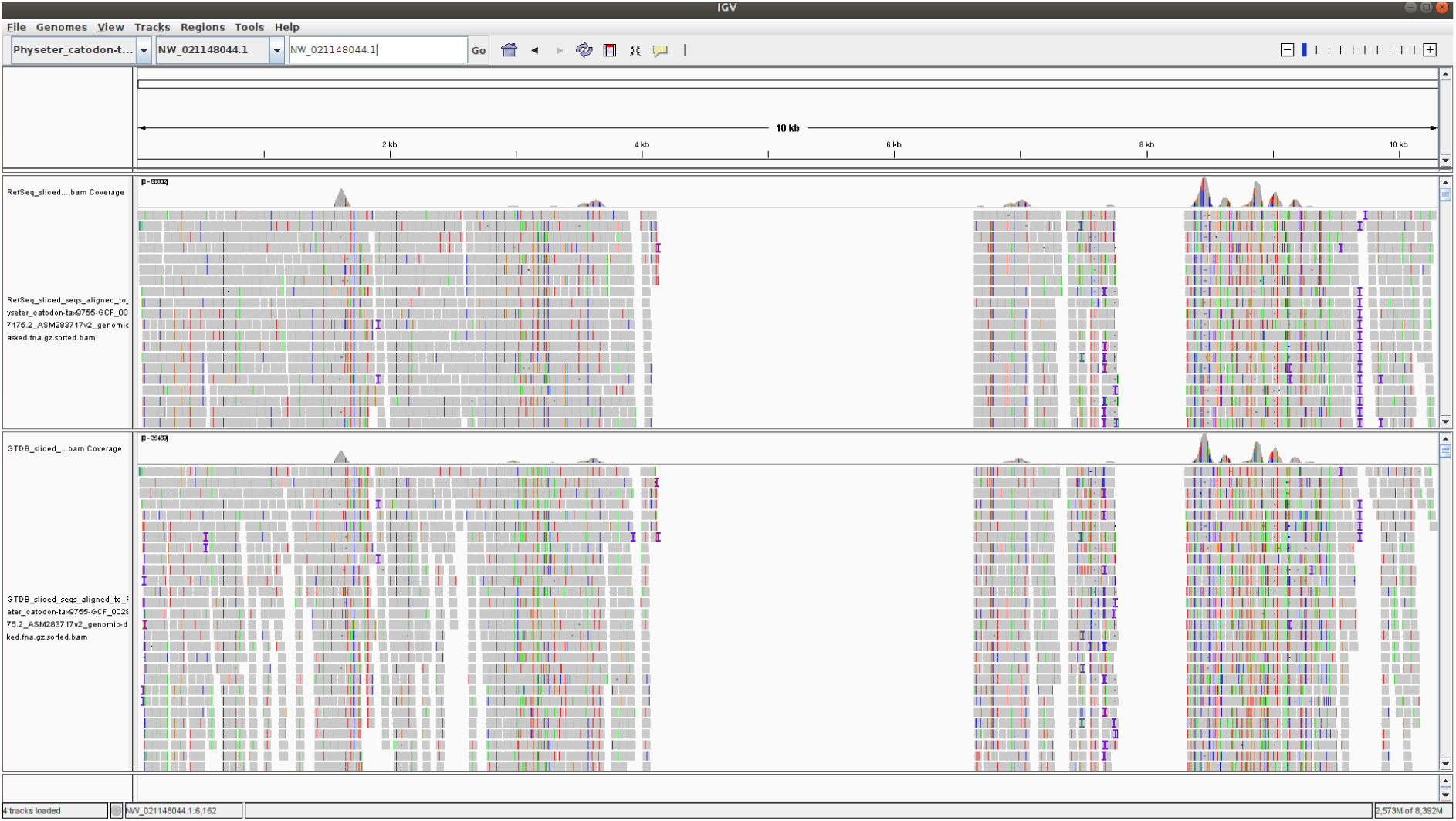
Comparison of coverage of a 10 kb region of the sperm whale (*Physeter catodon*, GCA_900411695.1) reference genome by microbial pseudo-reads produced from the microbial RefSeq (top) and microbial GTDB (bottom) databases. The visualization is performed using the Integrative Genome Viewer (IGV).

**Supplementary Figure 13.**
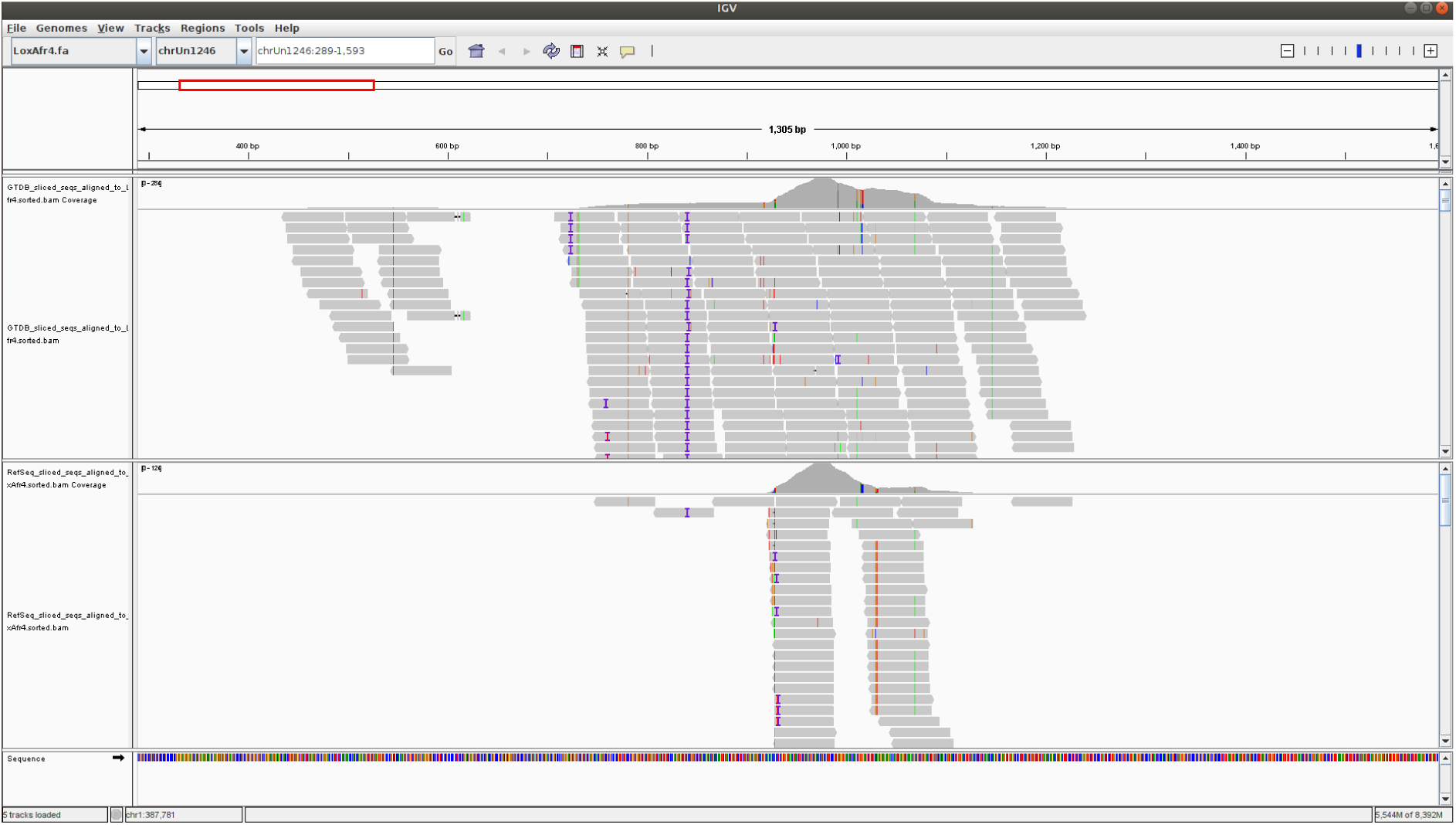
Comparison of coverage of a 1.3 kb region of the African bush elephant (*Loxodonta africana*, GCF_000001905.1) reference genome by microbial pseudo-reads produced from the microbial GTDB (top) and microbial RefSeq (bottom) databases. The visualization is performed using the Integrative Genome Viewer (IGV). The visualization demonstrates that microbial GTDB pseudo-reads are capable of discovering more microbial-like regions within the eukaryotic reference genome compared to microbial RefSeq pseudo-reads.

**Supplementary Figure 14.**
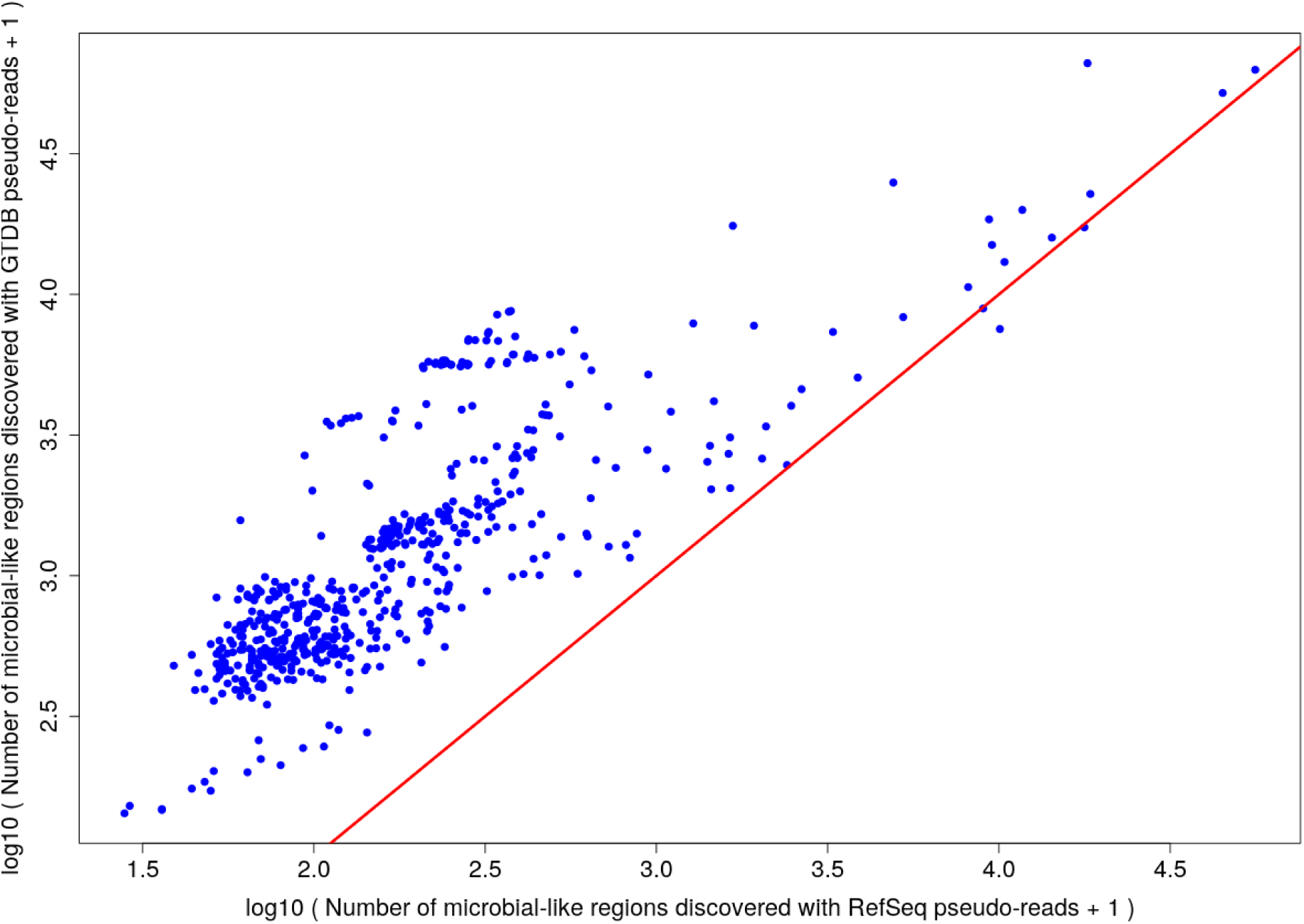
Comparison of numbers of microbial-like regions in mammalian reference genomes detected by using microbial GTDB and RefSeq pseudo-reads. One point represents one mammalian reference genome. Red diagonal line highlights equal counts for RefSeq and GTDB.

